# *In vivo* study of gene expression with an enhanced dual-color fluorescent transcriptional timer

**DOI:** 10.1101/564690

**Authors:** Li He, Richard Binari, Jiuhong Huang, Julia Falo-Sanjuan, Norbert Perrimon

## Abstract

Fluorescent transcriptional reporters are widely used as signaling reporters and biomarkers to monitor pathway activities and determine cell type identities. However, a large amount of dynamic information is lost due to the long half-life of the fluorescent proteins. To better detect dynamics, fluorescent transcriptional reporters can be destabilized to shorten their half-lives. However, applications of this approach *in vivo* are limited due to significant reduction of signal intensities. To overcome this limitation, we enhanced translation of a destabilized fluorescent protein and demonstrate the advantages of this approach by characterizing spatio-temporal changes of transcriptional activities in *Drosophila*. In addition, by combining a fast-folding destabilized fluorescent protein and a slow-folding long-lived fluorescent protein, we generated a dual-color transcriptional timer that provides spatio-temporal information about signaling pathway activities. Finally, we demonstrate the use of this transcriptional timer to identify new genes with dynamic expression patterns.

## Introduction

Changes in gene expression are one of the key mechanisms that organisms use during both development and homeostasis. Gene expression is a highly dynamic process, which not only bears critical information about regulatory mechanisms but also controls the fate of many biological processes^1,2^. For example, oscillatory or constant expression of the Notch effector *Hes1* dictates the choice of neuron stem cells between proliferation and differentiation^3^. In addition, defining the exact “on” and “off” timing of a relevant signal is vital to control different developmental events^4^. For example, during the development of fly compound eyes, simultaneous activation of EGF and Notch signals determines a cone cell fate^5^, while cells that experience sequential expression of EGF and the Notch-ligand Delta differentiate into photoreceptor cells^6^.

Documenting precisely the spatio-temporal changes in gene expression that occur in response to intrinsic and extrinsic signals is a challenging problem in cell and developmental biology. Traditionally, transcriptional reporters that drive expression of fluorescent proteins (FPs) under the control of signaling response elements (SREs) have been widely used to visualize the activities of transcriptional events; however, the slow degradation (half-life > 20 hr) of FPs makes it hard to achieve the temporal resolution needed to dissect the dynamic nature of gene expression. Recently, this problem has been addressed by the application of a fluorescent timer, a slow maturing fluorescent protein that changes its from blue to red in ∼7 hr^7^. Despite the still relatively long conversion time, this fluorescent timer has two additional limitations: the signal is hard to fix for long-term storage, and because it can be photoconverted from blue to red, this timer only allows one single image and prohibits live-imaging application^8^. Another strategy is the development of a destabilized version of GFP with a half-life of ∼2 hr, which is achieved by fusing GFP with a PEST peptide signal for protein degradation^9,10^. However, despite many *in vitro* successes, this strategy has met a major limitation when applied *in vivo* due to substantial loss of fluorescent intensity. Therefore, regular stable FPs are still the primarily choice for generating transcriptional reporters to study gene expression patterns *in vivo*.

Here, we address this problem by using translational enhancers to boost production of the destabilized reporters and demonstrate the advantages of using short-lived FPs to study dynamic gene expression *in vivo*. In addition, we generate a transcriptional timer that can be readily applied to study spatio-temporal activation of signaling pathways. Finally, we document how this transcriptional timer can be used, either using the UAS/Gal4 system or in an enhancer trap screen, to identify genes with dynamic expression.

## Results and Discussion

### Current limitations of stable transcriptional reporters and challenges of the destabilization strategy

Matching reporter dynamics with the activity of target genes is essential to faithfully recapitulate signaling activities^4^. Two primary kinetic properties dictate reporter activities: the “switch-on” and “switch-off” speeds. The “on” kinetic of FPs have been improved by engineering fast folding FPs, which shorten the maturation time of FPs from more than 1 hr to less than 10 min^11,12^. The “off” kinetics of FPs has been improved from more than 20 hr to around 2 hr by fusing FPs with PEST peptides that promote degradation^9^ (**Fig. 1a**). Although the advantages of using a short-lived reporter have been previously reported^9^, a systematic analysis of the differences between long-lived and short-lived reporters is still lacking. Using a protein synthesis and degradation model (**Fig. 1-figure supplement 1a-c**), we first simulated the dynamics of the reporters and demonstrated a significant improvement by decreasing the half-life (Tp_1/2_) of the reporters from 20 hr to 2 hr (**Fig. 1b-f**). Specifically, we illustrate this problem using simulated responses of FP reporters to four basic types of promoter activities: switch on, switch off, pulse activation, and oscillation. Compared to FPs with a half-life of 20 hr, FPs with a half-life of 2 hr have a 90% shorter response time (time to achieve 50% maximal intensity) during the “on” or “off” events, and up to four times larger dynamic range in the case of oscillatory expression (**Fig. 1 e,f**, **Figure 1-figure figure supplement 1a-c**).

**Fig. 1.**
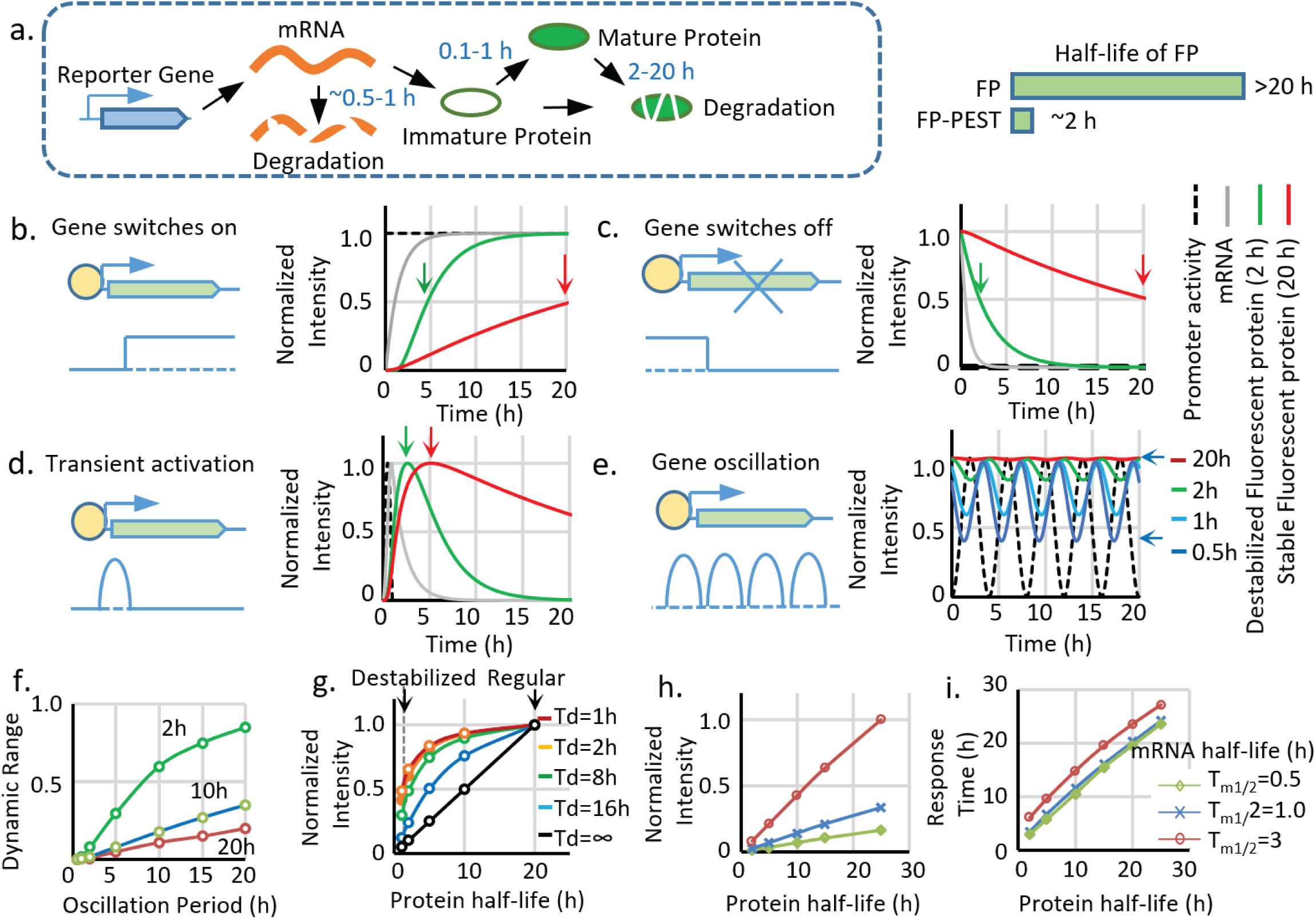
Advantages and limitations of destabilized fluorescent transcriptional reporters. **a.** Illustration of the biological processes that affect the final concentration of mature fluorescent protein, including transcription, translation, protein maturation, mRNA degradation, and protein degradation with the general half-life of mRNA, FP maturation time, and half-life of protein labeled. Destabiliation of the FP, achieved by fusion of FP with PEST domain, shortens the half-life of regular FP from over ∼20 hr to ∼2 hr. **b-d.** Comparison between simulated signals from a destabilized fluorescent reporter (green, *T*_*p1/2*_ =2 hr) and a regular fluorescent reporter (blue, *T*_*p1/2*_ =20 hr) following switch-on, switch-off, 1 hr pulse, and oscillation with a period of 2 hr. The half-life of mRNA (*T*_*m1/2*_) is set as 0.5 hr, and the protein maturation time (*τ*_*m1/2*_) is set as 0.1 hr. The time points when the fluorescent signals reach 50% of the maximal intensity (for switch-on and switch-off) and the maximum response (for transient pulse activation) are indicated by black arrows. **e.** Simulated signal intensities of fluorescent proteins with different protein half-lives to a sinusoid transcriptional oscillation with a period of 4 hr. The rest of the parameters are the same with above. The dynamic portion (the difference between the peak and valley) of the reporter (a half-life of 0.5 hr) is indicated by black arrows. **f.** The dynamic range (the difference between the peak and valley of the signal compared to its average intensity as indicated in **e.**) of reporters with indicated half-lives generated by sinusoid transcriptional activity with different length of the period. The dynamic range of the total signal positively correlates with the oscillation period and negatively correlates with the protein half-life. **g.** The intensity of the reporter signal for a constitutively active promoter (with no temporal variation) linearly depends on its protein half-life (black line). However, the maximal intensity of the reporter is less reduced by the shortened half-life for a shorter transient activation: a short-lived reporter (half-life of 2 h) shows a 90% reduction of maximal signal compared with a stable reporter (half-life of 20 h) for a constitutively active promoter. Nevertheless, the intensity is only reduced by 50% if the promoter is transiently activated for 1 hr. Td: the duration of a pulse transient promoter activation. **h**,**i.** Simulated changes in maximum fluorescent intensity and response time (time to reach 50% of the maximum intensity) of the reporters with different half-lives of protein and mRNA for a promoter switch-off event.

Although the destabilization strategy successfully leads to an FP with a shorter half-life, it is problematic as it causes significant loss of the signal (**Fig. 1 g-i**). Thus, at constant expression, the intensity of the maximum signal is linearly proportional to the protein half-life, and decreasing the half-life from 20 hr to 2 hr causes a 90% signal loss (**Fig. 1g**, **black curve**). The reduction of maximum signal intensity is also affected by different types of transcriptional activation. In the case of short pulsatile activation, a more transient activation (a shorter duration Td) is less sensitive to a reduction in the protein half-life (**Fig. 1g colored curves**); however, transient activation also triggers weaker reporter activity, which makes it more vulnerable to intensity reduction.

We further analyzed the effects of reducing mRNA half-life (Tm_1/2_) compared to protein destabilization (**Fig. 1 h, i**). Measurements of Tm_1/2_ of FPs in the literature are highly variable, ranging from several minutes to hours^13-15^, which is probably due to differences in the 3’ UTR used or different mRNA expression levels relative to the mRNA degradation machinery. According to our measurements (**Fig. 2, figure supplement 1e**), the Tm_1/2_ of FP reporters is about 0.5 hr. Therefore, we used 0.5 hr to 3 hr in our modeling. According to the model, the mRNA half-life significantly influences reporter intensity (**Fig. 1h**), which is approximately proportional to the Tp_1/2_* Tm_1/2_. In contrast, reducing mRNA half-life has much less effect on reporter response time, which is mainly controlled by Tp_1/2_ + Tm_1/2_ (**Fig. 1i**). Because in our system, Tp_1/2_ is much longer than Tm_1/2_, shortening the mRNA lifetime will significantly reduce signal intensity without endowing the reporter with more range in dynamic detection. Therefore, we decided to primarily use destabilized FPs in our study. For a system with a large Tm_1/2_ relative to Tp_1/2_, strategies to shorten mRNA lifetime by addition of RNA destabilizing sequence can be used^16^.

**Figure 2.**
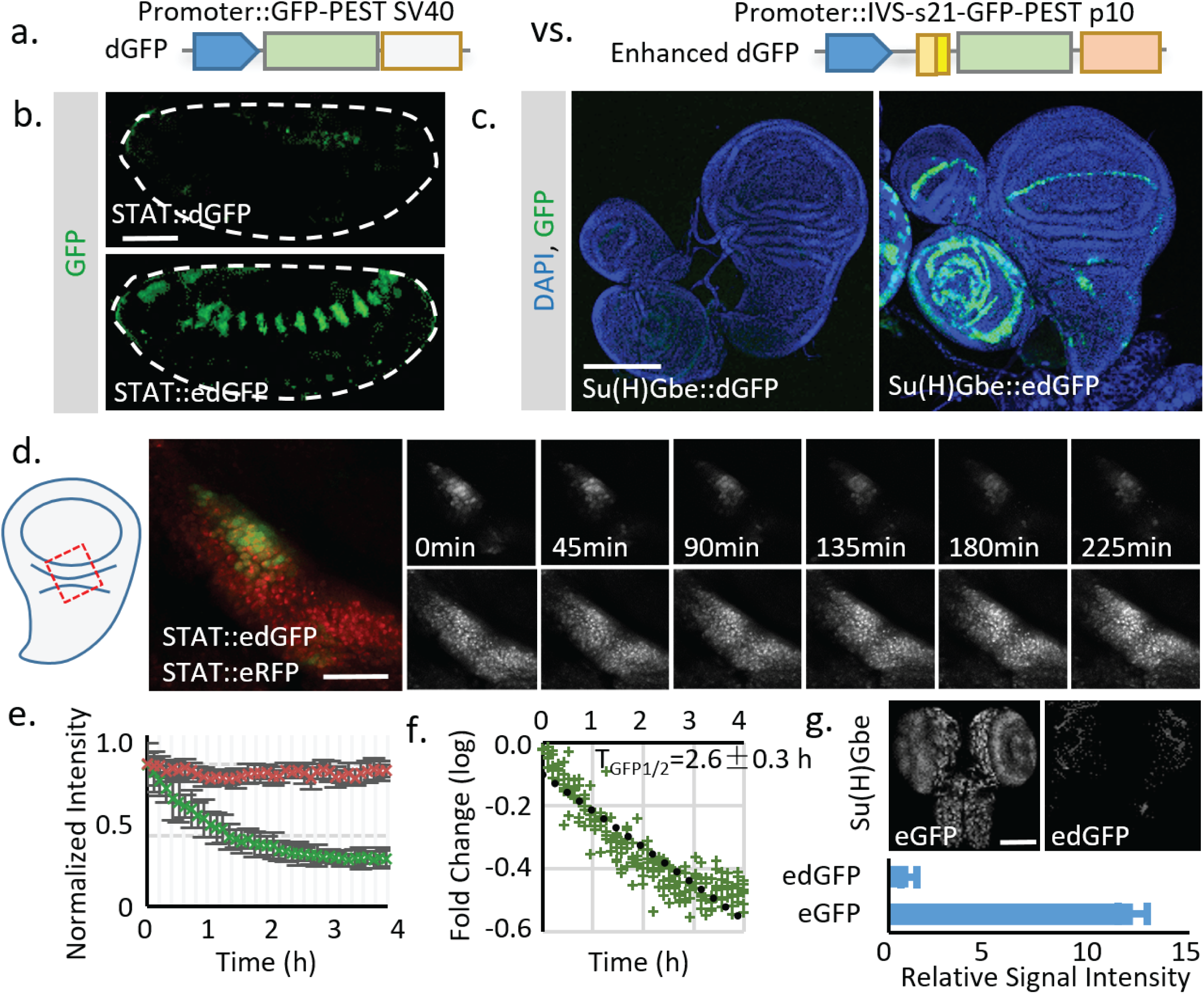
Using translational enhancers to increase the signal from destabilized fluorescent reporters for *in vivo* study. **a.** Illustration of the regular and destabilized GFP reporters. All the GFPs used in this study are the fast folding superfolder GFP (sfGFP). Destabilized GFP is labeled as dGFP, and dGFP reporter with translational enhancing elements is labeled as edGFP. All FPs used in this study contain the SV40 nuclear localization signal (NLS) at the N-terminus to facilitate signal segregation unless specified otherwise. **b**, **c.** Comparison of dGFP and edGFP controlled by 6XSTAT response element in fly embryos and Su(H)Gbe Notch responding element in 3^rd^ instar wing imaginal discs. Images were taken with identical exposure. The contour of the embryo is outlined. **d.** Dissected fly wing disc, expressing the *STAT∷edGFP* and *STAT∷ eRFP*, cultured *ex vivo*. Tissue was treated with 10 μM Actinomycin D to block transcription. STAT at the hinge region of the wing disc was imaged every 5 min for 4.5 hr. **e.** The intensities of both dGFP and RFP were measured over time. Data from 3 independent replicates were collected and plotted. **f.** The *in vivo* reporter half-life (*T*_*GFP1/2*_, representing effects of both *T*_*m1/2*_ and *T*_*p1/2*_) was estimated by linear regression of fluorescent intensity (in logarithmic scale). 95% confidence interval was calculated from linear regression. **g.** Regular GFP and dGFP are expressed under Su(H)Gbe together with translational enhancers. Images were taken under identical parameters. The total fluorescent intensity from both reporters was plotted below with the intensity normalized to the dGFP signal. Data were collected from 10 different brains for each genotype. Scale bar: **b.** 50 μm; **c**, **g.** 100 μm; **d.** 25 μm. Error bar: s.e.m.

### Application of translational enhancers to rescue the signal loss caused by destabilization

The significant loss of signal limits implementation of the destabilization strategy, especially in systems like *Drosophila* where a transgene is usually present in 1 or 2 copies per genome. To overcome this obstacle, we searched for ways to increase the signal of destabilized FPs. One possible solution is to use FPs with high intrinsic brightness. To test this, we used a fly codon-optimized sfGFP for its fast folding and bright fluorescence^11,17^. The effectiveness of destabilization was first tested in cultured fly S2 cells. Adding the PEST sequence from mouse ornithine decarboxylase (MODC) effectively reduced the half-life of sfGFP to ∼3 hr (**Fig. 2-figure supplement 1**). Next, to test the approach *in vivo*, we generated transgenic flies with destabilized sfGFP (dGFP) for two widely used signaling reporters: STAT (containing the STAT response element from *Socs36E*^18^) and Notch (containing the Notch response element Su(H)Gbe^19^). GFP signals were examined in tissues previously reported to show high STAT and Notch activities (embryo for STAT and wing imaginal disc for Notch). Destabilization reduced the signal intensity of these reporters to near background (**Fig. 2 a-c**). As further increasing the intrinsic brightness of FPs is challenging—even with the brightest FPs currently available the increase in signal intensity is still limited (less than two fold)^20^—we decided to test other strategies to increase the FP signal. One potential solution is to increase expression of the FP by expressing multiple tandem FPs^21,22^. However, this strategy makes cloning cumbersome and may render the insertion unstable due to recombination. Thus, we instead decided to use translational enhancing elements that have been demonstrated to increase protein production from mRNA by up to 20 fold^23,24^. These elements include a short 87-bp intervening sequence (IVS) from myosin heavy chain to facilitate mRNA export to the cytoplasm^24^, a synthetic AT-rich 21-bp sequence (s21) to promote translational initiation^23,25^, and a highly-efficient p10 polyadenylation (polyA) signal from baculovirus^26^. To test if these elements can be used to increase the reporter signals, we inserted the translational enhancing elements into reporter constructs containing dGFP (destabilized sfGFP) (**Fig. 2a-c**). Transgenic flies were generated by phiC31-mediated site-directed integration into the same genomic locus (attP40) to avoid potential position effect^27^. Strikingly, the addition of translational enhancers successfully increased the reporter signal with an expression pattern similar to that of previously reported stable reporters (**Fig. 2 a-c**)^19,28^. We further measured the *in vivo* half-life of the dGFP in live tissues by blocking transcription with Actinomycin (10 μM) and monitored degradation of the GFP (**Fig. 2d**). The result shows a reporter half-life (TGFP_1/2_ ∼2.6 ± 0.3 hr) similar to what was observed in cultured cells (∼3.7 ± 0.7 hr) (**Fig 2. d-f; figure 2-supplementary 1e**). We also tested the absolute signal intensity of regular GFP with the translational enhancer (eGFP) and destabilized GFP with the same enhancer (edGFP). The total signal intensity from edGFP is about 8% of what is observed for eGFP, consistent with a 91% reduction in protein half-life.

### Direct comparison between destabilized and stable reporters in live tissues

The effective increase of FP signal allows us to directly evaluate the activity of short-lived and long-lived traditional reporters. To achieve this, we generated a *STAT∷eRFP* reporter with the same translational enhancing elements and inserted it into the same locus (attP40) (**Fig. 3-supplementary 1a**). The activities of *STAT∷edGFP* and *STAT∷eRFP* reporters (as transheterozygotes) in developing embryos were examined under live imaging conditions (**Fig. 3a**). Compared to the stable RFP, the destabilized reporter showed a transient increase in STAT activity in tracheal pits (Tp), pharynx (Pr), proventriculus (Pv), posterior spiracles (Ps), and hindgut (Hg) (**Fig. 3a**, **figure 3-supplementary 1b**, **Supplementary Video 1**,**2**)^29^. To quantify the temporal changes of dGFP and RFP, the total fluorescent signals of both reporters were measured over time at the indicated region (arrowhead in **Fig. 3a**); the dGFP signal shows a definite improvement in response time (**Fig. 3b**). Using the half-lives estimated from the *in vivo* measurement (2.1 hr for dGFP and 18.5 hr for RFP) (**Fig. 3-figure figure supplement 1c-e**) and the reporter synthesis and degradation model (**Fig. 1-supplementary 1**), we further estimate the actual transcriptional activity of the reporter, which happens at an earlier stage (∼2 hr in advance of detectable dGFP reporter activity) (**Fig. 3b**). This result is consistent with the previously observed temporal expression of the STAT ligand *upd* from stage 9 to 12^30^.

**Figure 3.**
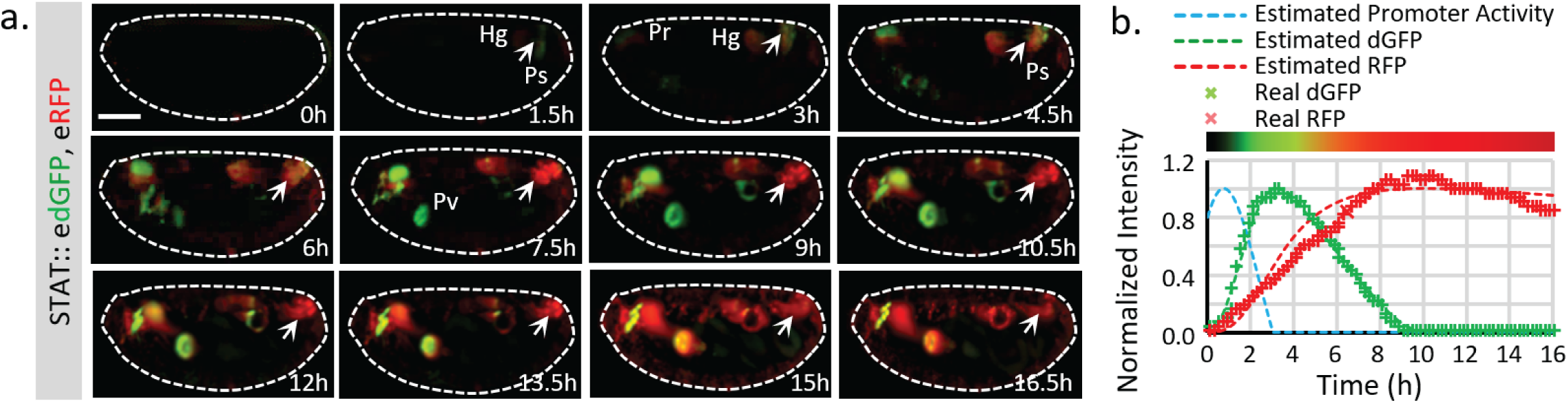
Live imaging of both destabilized sfGFP and stable RFP to reveal endogenous STAT dynamics. **b.** Time-lapse imaging of developing fly embryos expressing both *STAT∷edGFP* and *STAT∷eRFP* from stages 12 to 17 when STAT activity starts to increase. Maximum intensity z-projections of the mid-section (120 μm) are shown. Arrowheads indicate the posterior spiracles and hindgut region. The signal from the same structure turns from green to red over time. **c.** The total dGFP and RFP fluorescent signals within the posterior spiracles and hindgut region (indicated by arrows in **b**) are plotted as colored crosses. The estimated promoter activity was calculated using equations *(4)* (**Figure 1-figure Supplement, b**), and plotted as a dashed blue line. The simulated responses of dGFP and RFP from the estimated transcriptional signal using equations *(1)-(3)* (**Figure 1-figure Supplement, b**) are plotted as dashed green and red lines respectively. The merged intensity from estimated dGFP and RFP signals is illustrated as a color bar at the top of the plot. Scale bar: 100 μm

### Combining destabilized GFP and stable RFP to create a transcriptional timer

Short-lived reporters show clear advantages in revealing expression dynamics in live tissues. However, not all tissues are amenable to live imaging. Further, live imaging experiments are time-consuming and hard to adapt for large-scale studies. From our previous analyses, we noticed that dynamic information, including the initiation, maintenance, and reduction of transcriptional activities, can be directly estimated by comparing the ratio of fluorescence from stable vs destabilized reporters from a still image. Because the GFP matures faster than RFP (maturation time ∼0.1 h for sfGFP^31^ vs. ∼1.5 h for TagRFP^32^ used in this study), when the promoter switches on, a green signal is detected first. As the promoter remains active, both GFP and RFP signals reach a balance and such that their overlay produces a yellow color (when the max intensities of GFP and RFP are normalized). Furthermore, when transcription switches off, because the GFP signal decreases more quickly (the half-life of dGFP is ∼ 2 h, and the half-life of RFP is ∼ 20 h), only the RFP signal is left (**Fig. 4a**). Previously, ratiometric imaging of FPs based on their different properties, such as maturation time and sensitivity to specific ions, has been used successfully to measure protein life-time^31^, pH changes^33,34^, and Mg^2+^ concentration^35^. We reason that dynamic transcriptional activities might also be detected similarly using the ratio between dGFP and RFP.

**Figure 4.**
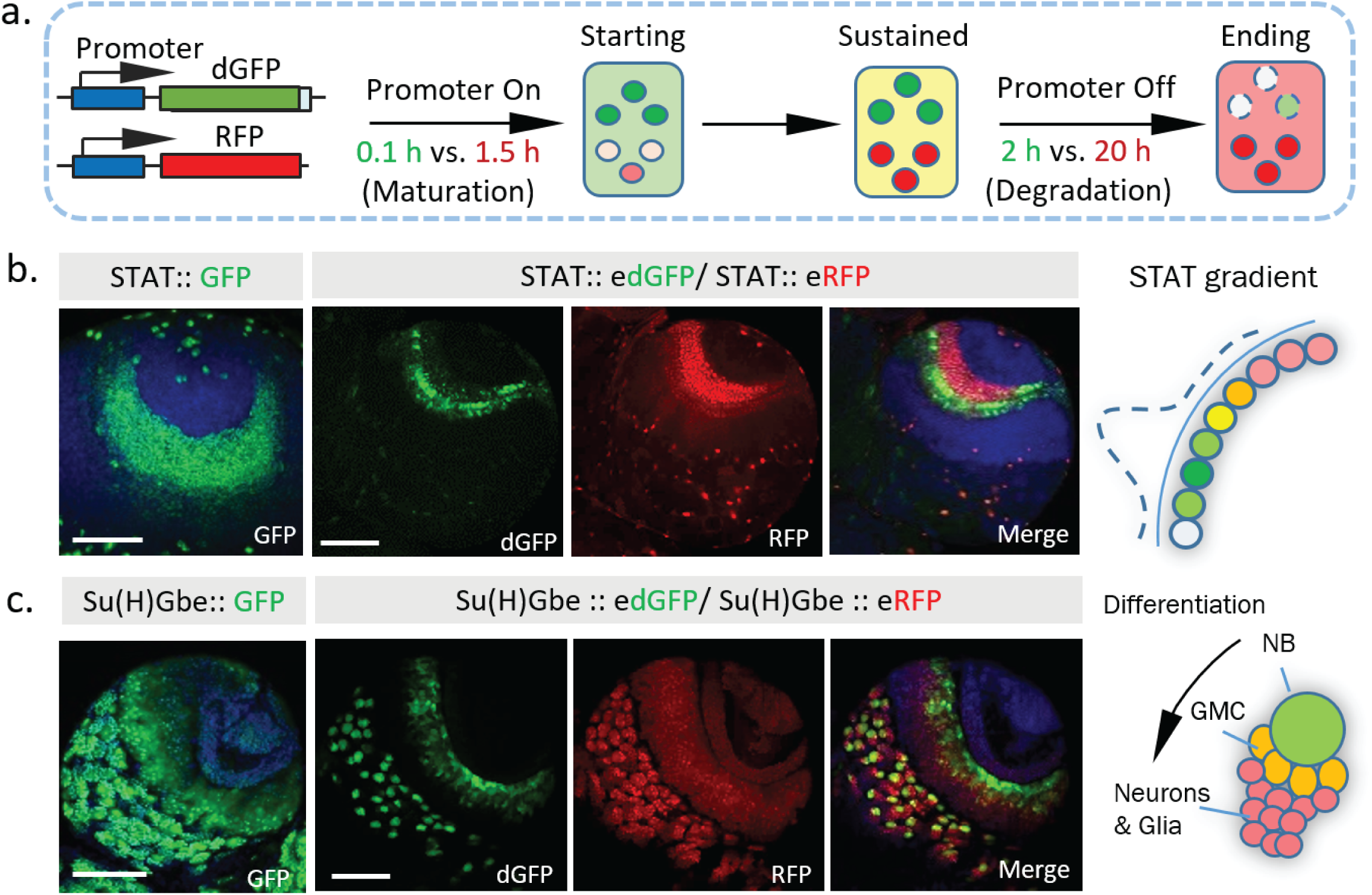
Dynamics of STAT and Notch activity revealed in fixed tissues. **a.** The destabilized GFP (dGFP) combined with stable RFP functions as a fluorescent timer to reveal transcriptional dynamics. The maturation and degradation half-lives of dGFP and RFP are indicated. **b. c.** A comparison between the regular GFP reporter with a combination between dGFP and RFP reporter under control of either STAT response elements or Notch response element Su(H)Gbe. Both reporters are visualized in 3^rd^ instar larval brains. NB, neuroblast; GMC, Ganglion mother cells. Scale bar: **a**,**b.** 50 μm

To test this strategy *in vivo* in a multicellular tissue, we examined STAT activity in the larval optic lobe. During development, STAT activity has been shown to act as a negative signal that antagonizes progression of the cell differentiation wave, which triggers the transition of neuroepithelial cells (NEs) into neuroblasts (NBs)^36^. A stable STAT reporter shows the spreading of STAT activity within the neuroepithelial region, similar to what we observe with *STAT∷eRFP* alone. In contrast, the signal from *STAT∷edGFP* together with *STAT∷eRFP* revealed a clear “green front” and “red rear” in the same region (**Fig. 4b**). This dynamic pattern, with a higher STAT activity at the boundary between NEs and NBs, is consistent with previous proposed wave-like STAT activity that propagates from the lateral to medial region ^36^. In addition, analysis of *STAT∷edGFP* and *STAT∷eRFP* in fixed larval optic lobes at different developmental stages further support this wave-like propagation model (**Fig. 4-figure Supplement 1a**).

Another example of the dynamic information revealed by combining edGFP and eRFP is the expression of the Notch reporter in larval brain NBs. NBs undergo an asymmetric cell division to generate smaller progeny (**Fig 4c**). Previous studies have shown that the Notch suppressor complex PON/Numb is preferentially localized to progeny cells^25, 26^ and that ectopic activation of Notch generates a brain tumor phenotype attributable to excess NBs, suggesting that Notch activity is required for NB self-renewal^37^. Strikingly, whereas the stable Notch reporter accumulates in both NBs and their progeny, the destabilized reporter is preferentially expressed in NBs (**Fig. 4c**, **Fig. 4-figure supplementary 1. b**), consistent with functional studies^37^. Combining the information from the edGFP and eRFP reporters reveals a clear Notch activity gradient that decreases as NBs differentiate. Importantly, in addition to its superior spatial resolution, the destabilized reporter also shows improved temporal resolution in NBs under live imaging conditions (**Fig. 4-figure supplementary 2**).

Our data suggest that combining edGFP and RFP creates a useful tool to study transcriptional dynamics, even in fixed samples. To facilitate this, we generated a multicistronic reporter containing both edGFP and RFP connected by the “self-cleaving” 2A peptide^38^, *edGFP-2A-RFP*, which we refer to as a transcriptional timer or “TransTimer” (**Fig. 4-figure supplement 3a**). Larval optic lobes expressing the transcriptional timer controlled by STAT response element show similar expression pattern as transheterozygous *STAT∷edGFP* and *STAT∷eRFP*, indicating that the multicistronic system is effective (**Fig. 4-figure supplement 3b**).

Destabilization of dGFP depends on protein degradation. Potential changes in the degradation speed of dGFP could affect the intensity of dGFP, which might distort our estimation of the real transcriptional activities. To test this possibility, we generated a TransTimer under the control of the fly Ubiquitin promoter (a constitutively active promoter in most fly tissues). *Ubi∷edGFP-2A-RFP* shows no significant variation in the green and red ratios in fly embryos or the larval brain (**Fig. 4-figure supplement 3c**). In addition, under control of other constitutive promoters, TransTimer also shows a relatively stable ratio between the two colors in different tissues (**Supplementary Table 2**), suggesting that changes in the FP ratios observed with TransTimer are primarily due to changes in transcriptional activity in the tested tissues, not cell type or tissue-specific differences in protein degradation. However, for specific organ or developmental stage, a control with a constitutive promoter for protein degradation changes is advisory.

### Generation of a transcriptional timer for use with the UAS/Gal4 system

Creating a TransTimer reporter for new signaling or target genes requires cloning of different signal response elements. Meanwhile, several large transgenic collections (up to several thousands) of Gal4 lines under the control of enhancers of different genes have been generated^39-41^. A TransTimer controlled by UAS would provide a quick way to test expression dynamics of Gal4 lines using a simple genetic cross (**Fig. 5a**). Thus, we generated *UAS∷TransTimer* transgenic flies and tested the UAS-controlled version in the adult *Drosophila* gut. TransTimer under control of *esg-Gal4*, a fly intestinal stem cells (ISCs) and enteroblasts (EBs) driver^42,43^, revealed particular cells within the stem cell group that turned “red,” distinguishing them from other “yellow” stem cells (**Fig. 5b**). Further analysis of these “red” cells showed that they down-regulate *esg* expression and up-regulate the enteroendocrine cell marker Prospero (Pros), suggesting that they are differentiating (**Fig. 5-figure supplement 1a**). In addition, the *esg*-controlled TransTimer shows significantly more dynamics in the developing intestine and regenerating intestine following Bleomycin treatment (a DNA damaging agent)^44^, consistent with the different level of activity of stem cells under these conditions (**Fig. 5b**, **Fig.5-figure supplement 1b**).

**Figure 5.**
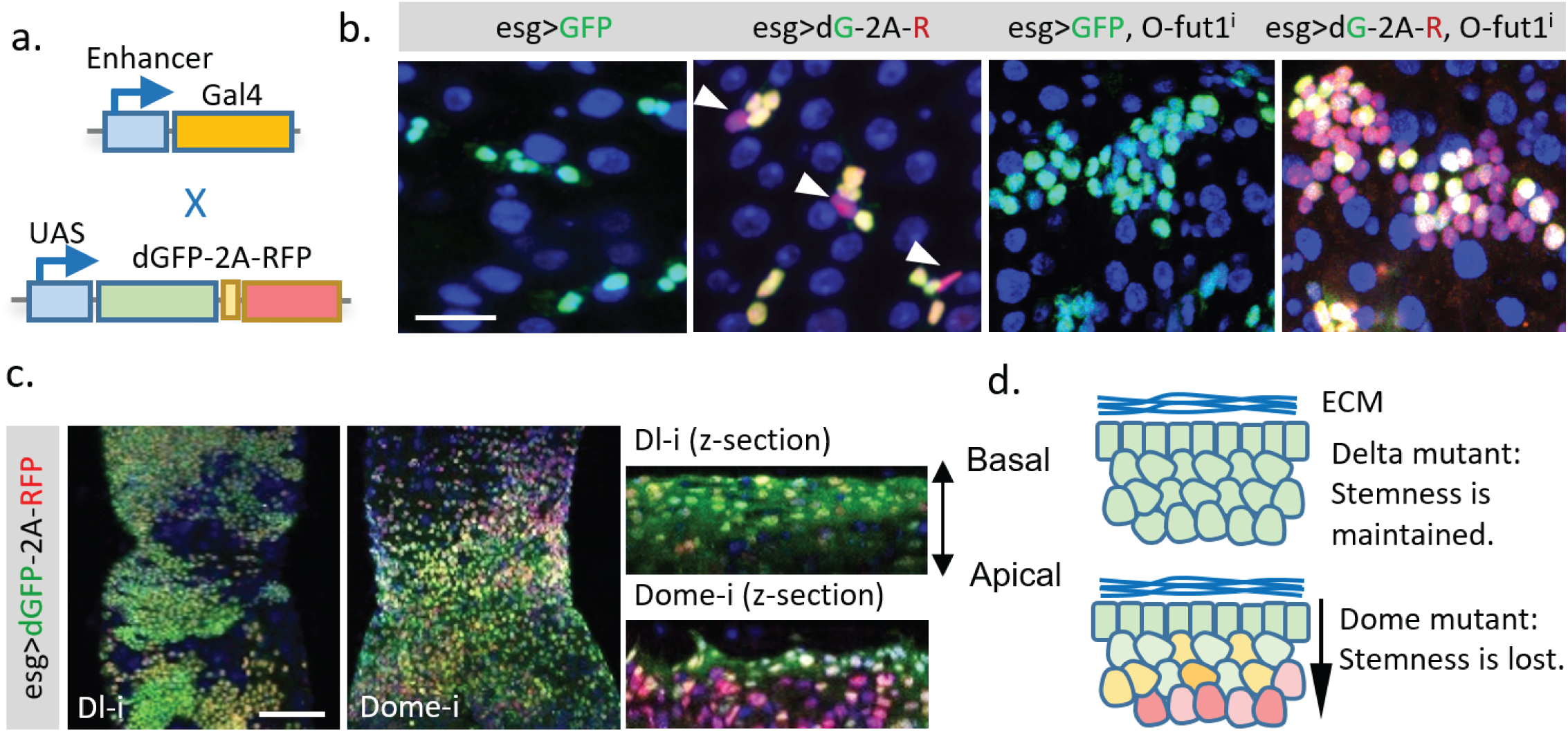
Application of the dGFP-P2A-RFP dynamic reporter to study gene expression dynamics of fly intestine stem cells. **a.** *UAS-TransTimer*, a UAS controlled multicistronic reporter containing dGFP and RFP connected by the P2A peptide was generated and crossed with the intestine stem cell driver *esg-Gal4*. **b.** Compared with the regular GFP reporter, *TransTimer* (*dGFP-P2A-RFP / dG-2A-R*) reveals the differentiating cells, which reduce the expression of the *escargot* (*esg*) stem cell marker (arrowheads). Knocking-down *O-fut1* inhibits Notch activity and causes ISC tumors. Compared to regular GFP, significant heterogeneity is revealed in the over-proliferating cell clusters using TransTimer. **c**,**d.** Intestine tumors are generated by knocking down the Notch ligand *Dl* or the cytokine receptor *Domeless* (*Dome*). The double-headed arrow indicates the z-direction with the basal side of intestine epithelium facing up and apical side facing down.

Next, we used TransTimer to examine the cell heterogeneity of different intestine tumors. In ISC tumors induced by knocking down *O-fut1*, an enzyme required for Notch maturation^42,43^, the regular GFP reporter showed only a moderate variation of the stem cell marker *esg.* By contrast, TransTimer revealed an evident decrease of dGFP compared to RFP in ∼70%-60% of the cells in the cluster, suggesting that a substantial heterogeneity in the tumor is caused by down-regulation of *esg* over time^45^ (**Fig 5b**). Tumors generated by knocking down either *Delta*, the ligand of Notch receptor, or *Domeless* (*Dome*), the transmembrane receptor of JAK/STAT signaling pathway, grow into a similar multilayered cell cluster (**Fig. 5c**)^46^. Interestingly, compared to *Dl* mutant tumors, where all multilayered cells maintain constant levels of *esg* (dGFP/RFP ratio), the inner layer of *Dome* mutant tumors shows clear reduction of *esg* expression (lower dGFP/RFP ratio) relative to the basal layer. This result suggests that *Dome* mutant tumors, unlike *Dl* mutant tumors, require direct contact with the basal membrane to keep their stemness (**Fig. 5d**).

### Application of TransTimer for the discovery of new genes with dynamic expression

As demonstrated above, *UAS-TransTimer* is an effective tool to discover expression changes when crossed with a Gal4 driver of interest. To further test the power of this approach to discover new genes with interesting expression dynamics, we screened ∼ 450 Gal4 lines using *UAS-TransTimer*^47^. 37 lines (∼8%) showed clear dynamic activities (substantial variation in dGFP/RFP ratio) in either larval brain, imaginal disc, or adult intestine (**Fig. 6a**, **Supplementary Table 1**), whereas the remaining Gal4s showed essentially uniform dGFP/RFP ratios, suggesting stable expression (images of representative control Gal4 lines are shown in **Supplementary Table 2**). Among the genes with dynamic expression patterns, we discovered the mechanosensitive channel *Piezo*, which is expressed in the posterior midgut specifically in EE precursor cells^48^. *TransTimer* driven by *Piezo-Gal4* **displays a spatially dynamic expression pattern** (separation between the “green” and “red” signals) (**Fig. 6b**). In addition, the “red” cells, which down-regulate Piezo expression, are positive for the EE cell marker Pros, consistent with the results of our previous study showing that Piezo+ cells differentiate into EE cells (**Fig. 6c**). In addition to *Piezo*, we also identified new uncharacterized genes with dynamic expression patterns in a subpopulation of *esg+* cells (**Fig. 6d**, **Fig. 6e**). Further studies of these genes will be required to determine whether they are markers of partially differentiated cells like Piezo or if their expression levels oscillate in the stem cells.

**Figure 6.**
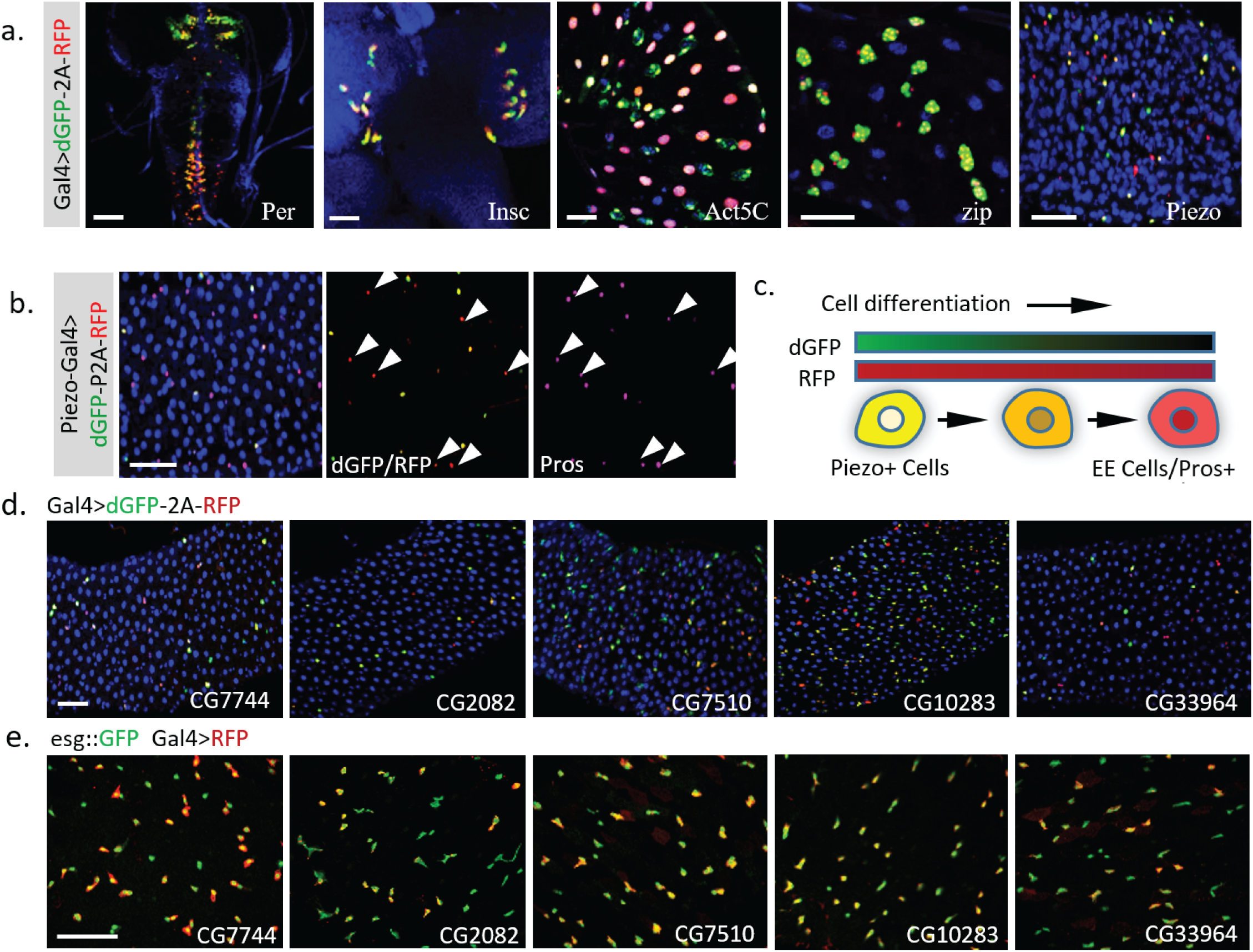
Dynamic pattern of TransTimer driven by different Gal4 drivers. **a.** Examples of Gal4 lines that show clear dynamic patterns with TransTimer (*UAS-dGFP-P2A-RFP*) in various organs: *Per-Gal4*, controlled by the circadian rhythm regulator Period (larval brain and ventral ganglion); *insc-Gal4*, expressed in Type II neuroblasts (larval brain); *Act5C-Gal4*, controlled by the *act5C* enhancer (larval intestine); *GMR51F08-Gal4*, controlled by the enhancer from the fly myosin heavy chain *Zipper* (larval intestine, adult intestine precursor cells); *Piezo-Gal4 (BL58771)*, controlled by the cloned enhancer from mechanical sensitive ion channel *Piezo* (adult intestine). **b.** Dynamic reporter under control of *Piezo-Gal4[KI], a Gal4* knock-in after the first ATG of *Piezo*. Piezo+ cells with high RFP and low GFP are positive for the EE cell marker Prospero (Pros). **c.** Illustration of the differentiation process from Piezo+ EP (enteroendocrine precursor) to Pros+ EE cells. **d.** The expression dynamics of *UAS-TransTimer* driven by different Gal4 lines in the fly intestine. **e.** Gal4 activities are detected in a subpopulation of the *esg*+ stem cells. Stem cells are marked by *esg∷GFP* and the expression of Gal4s are revealed by *UAS-RFP*. Scale bar: **a-e.** 50 μm.

We also generated transgenic flies with TransTimer controlled by a minimal promoter that is silent unless activated by nearby enhancers and randomly mobilized the transgene in the fly genome to identify endogenous enhancers with dynamic activities (**Fig. 7a**, **Fig. 7-figure supplement 1**). After screening ∼400 independent enhancer trap lines, we identified 46 unique lines that showed fluorescent signals in the larval brain, imaginal disc, or adult intestine. 17 of these 46 lines show clear expression dynamics, suggesting that TransTimer can detect expression changes at endogenous levels (**Fig 7b**, **Fig. 7-figure supplement 1b**, **Supplementary Table 3**). To validate the screen, we tested the expression and function of new genes identified in this enhancer trap screen. Since we are particularly interested in new lines that show exclusive expression in stem cells, we chose TransTimer insertions near the promoters of *Tsp42Ea*, a Tetraspanin protein, and *CG30159*, an evolutionarily conserved gene with unknown function - as the function of these genes had yet not been characterized in fly intestine. A Gal4 line (*NP1176-Gal4*), located closely (within 250 bp) to the TransTimer insertion site at the promoter of *Tsp42Ea* and *CG30159*, also shows specific expression in both larval and adult intestine stem cells (**Fig. 7d**), which is very similar with the expression pattern revealed by TransTimer (**Fig. 7b**). Knocking down *CG30159* significantly reduces stem cell numbers, suggesting that *CG30159* is required for maintenance of intestinal stem cell (**Fig. 7e**,**f**). The human homolog of *CG30159* is *C3orf33*, which has been identified as a regulator of the extracellular signal-regulated kinase (ERK) and predicted to be a secreted peptide due to the presence of signal peptide at its N-terminus^49^. Its function in intestinal stem cells requires further investigation.

**Figure 7.**
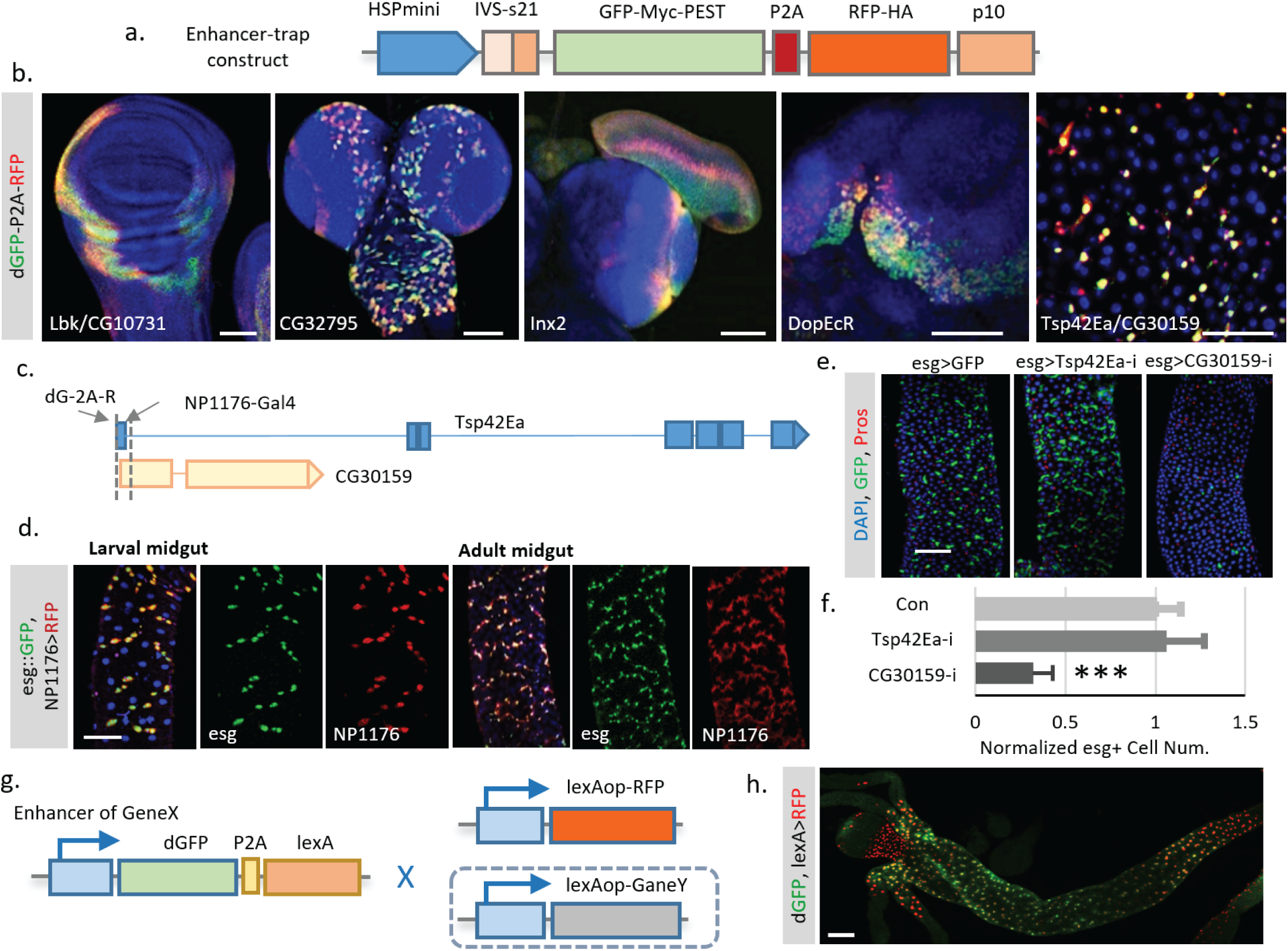
Enhancer trap screen for endogenous transcription dynamics. **a.** A P-element containing *dGFP-P2A-RFP* (TransTimer) was randomly mobilized in the fly genome to identify enhancers with dynamic expression. edGFP and RFP are tagged by Myc and HA epitopes, respectively, to allow signal enhancement by immunohistochemistry. **b.** Examples of enhancer trap lines that show dynamic patterns in different organs: *Lbk*, an Immunoglobulin-like protein, or *CG10731*, the subunit S of mitochondrial ATP synthase complex (wing disc); *CG32795*, a novel membrane protein (larval brain); *inx2*, a gap junction protein (larval brain and eye disc); *DopEcR*, Dopamine and Ecdysteroid receptor (larval brain); and *Tsp42E*, a tetraspanin protein; and *CG30159*, a novel gene with unknown function (adult intestine). **c.** The *dGFP-P2A-RFP* and the *NP1176-Gal4* enhancer trap insertions at the promoter region of *Tsp42Ea* and *CG30159*. Exons are shown in the diagram (blue for *Tsp42Ea*, yellow for *CG30159*). *Tsp42Ea* and *CG30159* share the same promoter region. **d.** *NP1176-Gal4* (marked by *UAS-RFP*) shows stem cell expression like *esg* (marked by *esg∷GFP*). **e**, **f.** Knocking down *CG30159* significantly reduces stem cell numbers (*esg*+ cells) in the intestine, while RNAi against *Tsp42Ea* does not have a significant phenotype. Number of *esg*+ cells were quantified within 100 μm^2 regions from n=8 (*GFP* control), n=7 (*Tsp42Ea-i*), and n=9 (*GC30159-i*) adult fly intestines. P-value <0.001. Cell numbers are normalized according to the control. **g.** Schematic of the TransTimerLex for enhancer trap with RFP replaced by the bacterial transcriptional repressor LexA. This reporter allows both two-color contrast and further genetic manipulation of the target cell population using the LexA/lexAop binary expression system. **h.** An enhancer trap line, under potential control of *Larp* enhancer, was crossed with *lexAop∷RFP*. The anterior region of the larval intestine is shown, revealing a decrease of transcriptional activity in the proventriculus. Scale bar: Scale bar: **b**, **h.** 50 μm**; d**, **e.** 100 μm.

As we have shown above, the enhancer trap screen with TransTimer can effectively detect expression dynamics *in vivo*. However, this screen can only detect gene expression and cannot be used to manipulate the target cell population. To extend the application of TransTimer, we replaced the RFP with lexA, a yeast transcriptional factor used as a binary expression system together with its binding sequence lexA operator (lexAop)^50^. This *dGFP-P2A-lexA* construct can not only detect expression dynamics when crossed with lexAop-RFP but also manipulate gene expression in labeled cells in the presence of an additional lexAOP-controlled transgene (**Fig. 7g**). We refer to *dGFP-P2A-lexA* as “TransTimerLex”. To test the feasibility of this strategy, we generated transgenic flies containing the TransTimerLex insertion and randomly mobilized the element in the fly genome. From our pilot screen (∼20 independent lines), we identified one insertion under control of Larp, a transcriptional factor, which shows clear expression dynamics in the larval intestine (**Fig. 7h**). This result suggests that the TransTimeLex system will be a useful way to both identify new genes and manipulate gene expression in the corresponding cells.

### Concluding Remarks

In this study, we described a general and straightforward strategy to use destabilized transcriptional reporters *in vivo* and demonstrated its power in revealing the spatio-temporal dynamics of gene expression, which is missed by conventional transcriptional reporters. In addition, we generated a dual-color TransTimer that encodes the transcriptional dynamics into a green-to-red color ratio which can be analyzed in fixed tissues. This TransTimer provides a unique opportunity for large-scale screens for *in vivo* expression dynamics in all types of tissues. Further, our study indicates that TransTimer is effective for the discovery of new genes with interesting expression patterns, either using a candidate gene approach or random genome-wide screening. Although we only tested this strategy in *Drosophila*, there is no major technical limitation to adapt it to other organisms. Therefore, we expect that this new method will widely facilitate studies in *Drosophila* and other organisms.

## ACKNOWLEDGMENTS

We thank Douglas Richardson at Harvard Biological Imaging Center for technical support and advice, and Ben Ewen-Campen, Benjamin Housden, Stephanie Mohr, and Muhammad Ahmad for comments on the manuscript. This work was supported by the Damon Runyon Cancer Research Foundation (L.H.) and NIH (R21DA039582). N.P. is an investigator of the Howard Hughes Medical Institute.

## CONTRIBUTIONS

The project was conceived and designed by L.H. and N.P. L.H., J.H., and J.F. performed the construction, *in vitro*, and *in vivo* characterization of destabilized fluorescent reporters. L.H. performed the live-cell imaging experiments and analyzed the data. R.B. and L.H. performed the enhancer screen for dynamic expression patterns. L.H. and N.P. wrote the manuscript with input from all of the authors.

## Materials and methods

### Molecular Biology

cDNAs of sfGFP and TagRFP, codon optimized for *Drosophila*, were a kind gift from Dr. Hugo Bellen^17^. HSPmini-IVS-p21 and p10 were amplified by PCR from pJFRC81 (Addgene)^23^. MODC sequence was from pBPhsFlp2∷PEST vector (Addgene)^51^. The vector containing LHG, which encodes a chimeric protein consisting of the LexA DNA binding domain and the Gal4 transcriptional activation domain, was a kind gift from Dr. Konrad Basler^50^. pUAST4 and pWALIUM10-roe were from our lab stock collection. Primers and gBlocks were obtained from Integrated DNA Technologies. PCR was performed with the proofreading enzyme Phusion (NEB). Plasmid purification, PCR purification, and gel extraction were performed with QIAprep Spin Miniprep Kit (QIAGEN), QIAquick PCR Purification, and QIAquick Gel Extraction Kits (QIAGEN), respectively. In-fusion cloning and Gateway cloning were performed using In-Fusion HD Liquid Kits (Clontech), and BP and LR Clonase Enzyme Mixes (ThermoFisher Scientific). All cloning experiments were verified by DNA sequencing. Cloning details and DNA sequences are shown in **Supplementary Data 1**.

#### *Drosophila* Genetics

The following fly lines were obtained from the Bloomington *Drosophila* Stock Center: *Delta2-3*(99B) (BL3629), *Per-Gal4 (BL7127), Trx-Gal4 (BL40366), Ogre-Gal4 (BL49340), RhoGAP-Gal4 (BL45252), Gcm-gal4 (BL35541), dMyc-Gal4 (BL47844), Hh (BL49437), Antp (BL26817), Act5C (BL9431), BL48187, dMyc(BL47844), piezo-Gal4 (BL58771, BL59266), ppk-Gal4 (BL32078, BL32079), Ubi-Gal4 (BL32551), MESK2-Gal4 (BL67434), CG7744-Gal4 (BL76662), CG2082-Gal4 (BL76181), CG7510-Gal4 (BL66861), CG10283-Gal4 (BL76152), CG33964-Gal4 (BL76742)*, and *dMef2-Gal4 (BL27390). NP1176-Gal4* was from DGRC (Kyoto Stock Center)*. GMR-Gal4, Da-Gal4, tubGal4*, and *esg-Gal4* were from lab stocks. *Insc-Gal4, ase-Gal80ts* was a gift from Dr. Dong Yan^52^ and *Dl-Gal4* from Dr. Steve Hou^53^. All flies were maintained on cornmeal-yeast-agar media. Stocks were kept at room temperature with a 12/12 light/dark cycle.

#### *Drosophila* S2R+ cell culture and Western blotting

*Drosophila* S2R+ cells were grown in Schneider’s *Drosophila* Medium (SDM) (Invitrogen) containing 10% heat-inactivated fetal bovine serum (FBS) at 25°C. Sub-confluent S2R+ cells were seeded in 6-well plates and subsequently transfected using Effectene Transfection Reagent (QIAGEN). Cells were cultured for 48 hr before experiments. 10 μM (final concentration) Actinomycin D was used to block RNA synthesis, and 100 μg/ml (final concentration) cycloheximide was used to block protein synthesis. Cells were treated with the indicated drugs up to 4.5 hr before significant cell death was observed. Plasmids expressing pUbi-dGFP-Myc (0.03 ug), and pUbi-RFP-HA (0.01 ug), together with empty plasmids (to reach a total of 0.3 ug of DNA) were added in each 6-well plate during transfection. The dilution of the expression plasmid was important: we observed that too much protein expression saturates the degradation machinery and prolongs the observed half-life. S2R+ cells were harvested by centrifugation and lysed in RIPA buffer. Proteins were separated on a 10% SDS-PAGE gel and analyzed by Western blotting. Quantitative Western blots were performed as previously described^54^. Images were acquired using a LI-COR Odyssey Classic imager and analyzed using NIH ImageJ.

### Generation and test of enhancer trap lines using dynamic enhancers

Dynamic reporters were integrated into the fly genome using P-element mediated transformation by injection into *w1118* embryos^55^. Transgenic lines were balanced and mapped using *w*; Sco/CyO; MKRS/TM6B*. Then, 6 independent lines were generated by crossing with *w*;Sp/CyO; Sb,P(Delta2-3)99B/TM6B,Tb+* (BL3629). Males from the F1 generation with red eyes and carrying the *CyO* balancer were further crossed individually with *w1118* females. F2 males with red eyes that co-segregate with the *CyO* balancer were used in the initial screen.

Detailed crosses for the enhancer screen are shown in **Supplementary Fig. 6a**. ∼400 fly lines were recovered from the F2 generation. 3rd instar larval brains, imaginal discs, and adult intestines of these flies were dissected and examined for GFP and RFP signal using a Zeiss LSM 780 confocal microscope. P-element insertion sites were mapped by Splinkerette PCR^56^. PCR primers specific for 5 and 3 prime ends of P-elements were used as previously described^57^. Genomic sequences flanking the P-element insertion sites were recovered and shown in **Supplementary Data 2**. These sequences were used in BLAST searches against the *Drosophila* Genome Database.

### Immunostaining

Immunostainings of *Drosophila* intestines were performed as previously described^42^. The following antibodies were used: mouse anti-Prospero (1:50, Developmental Studies Hybridoma Bank), mouse anti-HA (1:500, Abcam, ab18181), rabbit anti-Myc tag (1/250, Cell signaling), goat anti-mouse IgGs conjugated to Alexa 647 (1:500, Molecular Probes), mouse IgGs conjugated to Alexa 488 (1:500, Molecular Probes), IRDye® 800CW Goat anti-Rabbit IgG (1:10,000 LI-COR P/N 926-32231), and IRDye 680RD Goat anti-Mouse IgG (1:20,000 LI-COR P/N 926-68070). Dissected fly tissues were mounted in Vectashield with DAPI (Vector Laboratories). In all micrographs, the blue signal shows the nuclear marker DAPI. Fluorescence micrographs were acquired with a Zeiss LSM 780 confocal microscope. All images were adjusted and assembled in NIH ImageJ.

### Modeling of FP production, maturation, and degradation

The model used to calculate reporter synthesis, maturation, and degradation was modified from previously described equations^58^ with the addition of an equation for mRNA degradation. Briefly, degradation of mRNA and the synthesis rate of premature (nonfluorescent) protein (NP) is proportional to the mRNA concentration (R). Generation of the mature reporter (MP), modeled as a first-order chemical reaction, only depends on the concentration of NP. Protein degradation is modeled independently of the maturation process. The degradation rates of mRNA and proteins are first modeled based on Michaelis-Menten (MM) function **(Fig. 1-figure supplement b, equations 1-3)**, which considers the potential saturation of the degradation machinery. When the substrate concentration is significantly smaller than the Michaelis constant *Km*, the equations can be simplified with the half-life of the mRNA and protein explicitly displayed **(Fig. 1-figure supplement b, equations 1’-3’)**. Dilution by cell division is not included in this model because the fluorescent signal is analyzed in a cell cluster rather than in individual cells, and cell division does not affect the total intensity from the entire cell group and no significant changes in degradation speed have been observed between different cells (**Fig. 3 figure supplement 1**). With this first-order kinetic model, the transcriptional activity of the promoter *F(x)* can be calculated through equation from the observed fluorescent reporter signal *[MP]* (4). For sfGFP, the maturation time is ∼0.1 hr, which is much smaller than its protein half-life, such that equation **4** can be further simplified as **4’**. To calculate *F(x)*, the dGFP signal was fitted with a polynomial function (order=4) to generate the first and second derivatives.

### Live-cell imaging and data analysis

Live-cell imaging of developing embryos and dissected larval brains was performed as previously described^59-61^. Images were captured on a Zeiss Lightsheet Z1 microscope using a 20X (N.A. 1.0) lens. A z-stack of the dual-color image (488 nm excitation/500-550 nm detection for GFP, and 561 nm excitation/580-650 nm detection for RFP) was recorded at 10 min intervals. This interval was chosen empirically to minimize photobleaching without losing temporal information. Photobleaching was measured by continuously imaging of the sample for 50 frames for 10 min and adjusted during image processing. Images of fixed tissue were captured on a Zeiss LSM 780 confocal microscope. Total fluorescent intensity in 3D volume was acquired using Imaris image analysis software (Bitplane). The rest of the analysis was completed using NIH ImageJ with customized macros. Simulation of the model was completed in MATLAB. The Student’s unpaired, two-tailed t-test was used to determine statistical significance between samples.

**Figure 1-figure Supplement.**
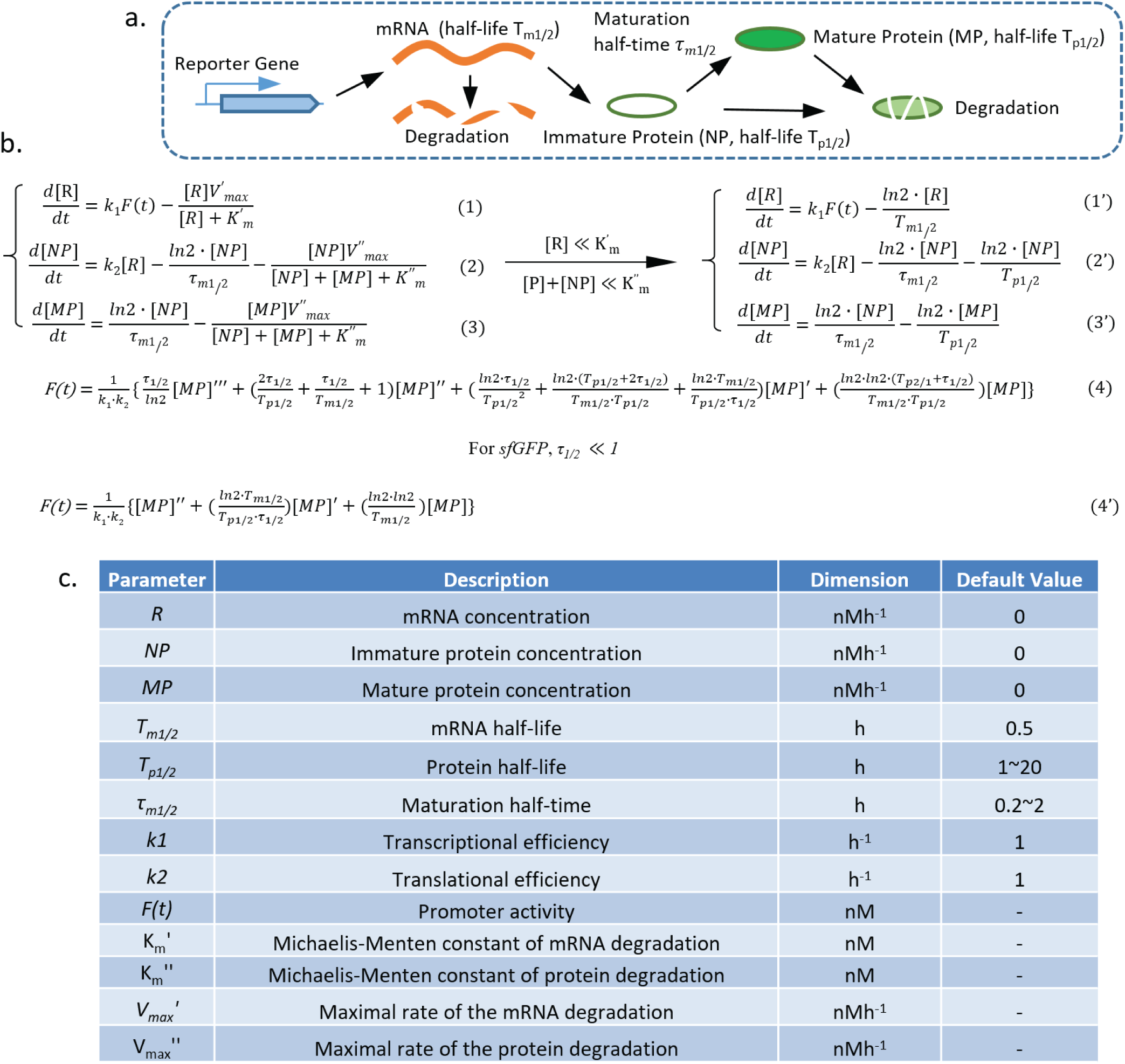
Model of the transcriptional reporter synthesis, maturation, and degradation. **a.** Illustration of the biological processes that affect the concentration of mature fluorescent protein, including transcription, translation, protein maturation, mRNA degradation, and protein degradation with key parameters labeled. **b**,**c.** Equations describing the model generated by following the law of mass action, with the parameters shown in the table below.

**Figure 2-figure Supplement 1.**
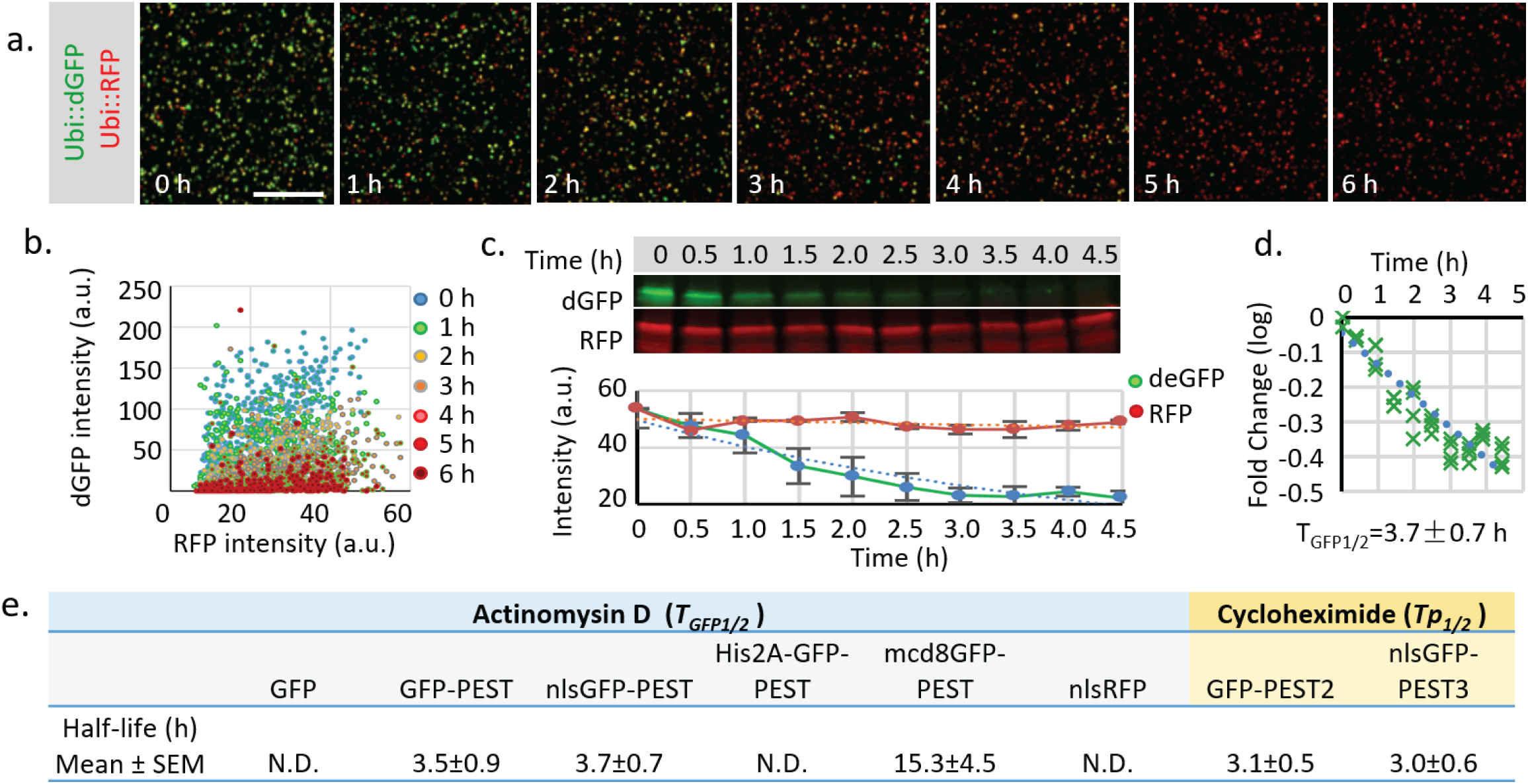
Measurement of the half-life of destabilized GFP in cultured *Drosophila* S2 cells. **a.** *Drosophila* S2 cells transiently transfected with both *Ubi∷nls-sfGFP-Myc-PEST* and *Ubi∷nls-RFP-HA* were treated with 10 μM Actinomycin D to block RNA synthesis for 6 hours. Images with identical exposure time were taken every hour after drug addiction. Scale bar: 100 μm. **b.** The plot of fluorescence intensity of dGFP and RFP in cultured cells at different time points. **c.** S2 cells treated with the transcriptional blocker Actinomycin D were harvested every 0.5 hr from time 0.0 hr to time 4.5 hr. Samples were analyzed by fluorescent western blot using rabbit anti-Myc (800 nm, Green) and mouse anti-HA (700 nm, Red). Quantification of fluorescent intensity from 3 independent experiments. Error bar: s.e.m. **d.** Fold change of fluorescent intensity was plotted over time (logarithmic scale). The combined half-life of both mRNA and protein (*T*_*GFP1/2*_) was calculated by linear regression using least squares estimation. The expected value and 95% confident interval of *T*_*GFP1/2*_ was calculated from the linear regression. **e.** Table of measured half-lives of fluorescent reporters (*T*_*GFP1/2*_, including both mRNA and protein half-life) measured using western blot after Actinomycin D treatment (in blue cells). Protein half-life (T_p1/2_) was measured using western blot after cycloheximide (100 μg/ml) treatment to block protein synthesis (in yellow cells). Regular GFP, RFP or GFP-PEST fused with His2A is too stable to be reliably estimated within the 5 hr treatment period.

**Figure 3-figure supplement 1.**
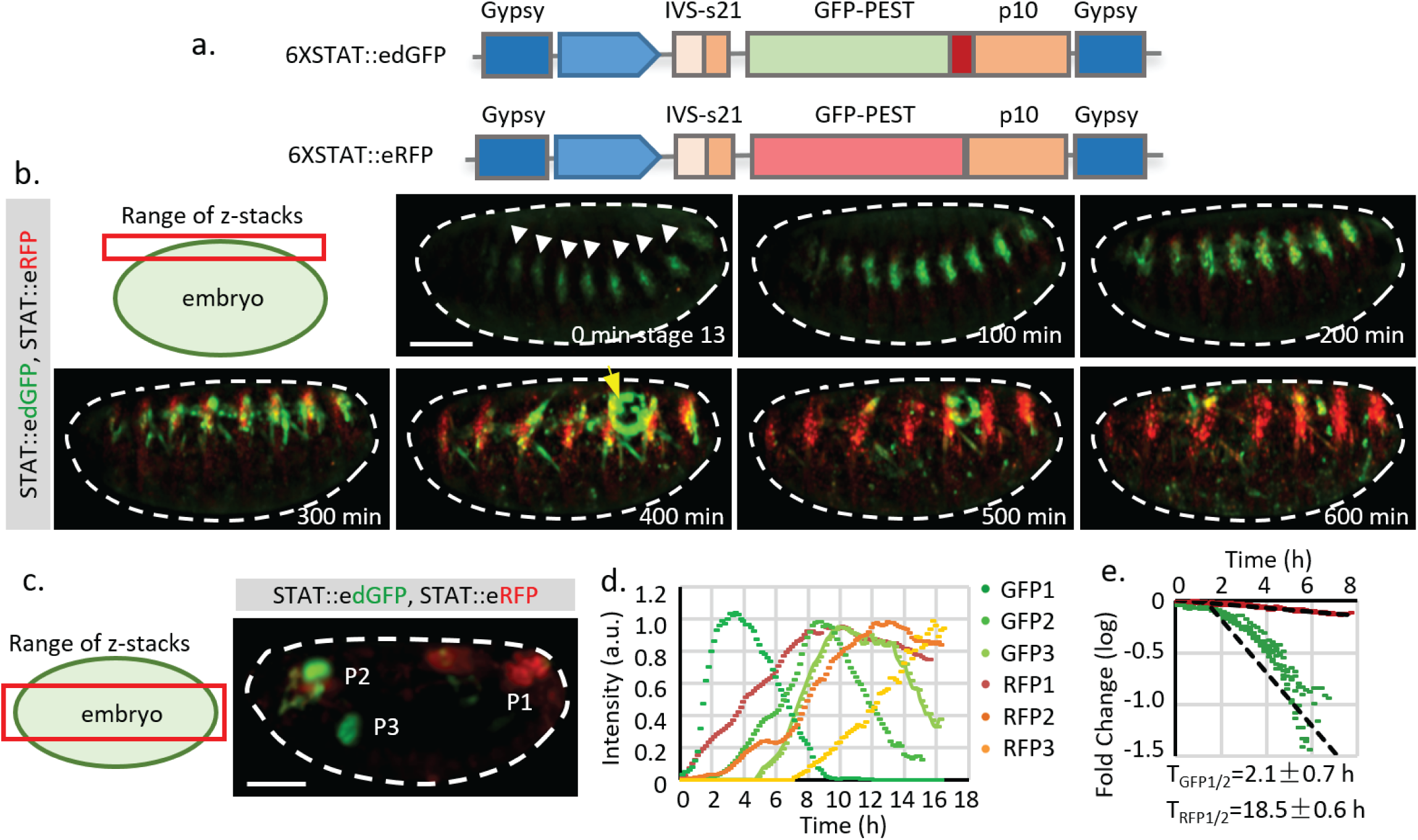
Quantification of STAT reporter dynamics. **a.** The STAT reporter consists of gypsy insulators at each end to prevent potential influence of nearby promoters, a *6XSTAT* response element from STAT effector *Socs36E*, a minimal promoter from the *hsp70* heat shock gene (HSPmin), an intervening sequence (IVS), a Syn21 sequence, a fly codon-optimized fluorescent protein with nuclear localization signal (NLS), a PEST sequence (to destabilize the GFP), and p10 polyA signal. **b.** Frames from live imaging of *STAT∷edGFP* and *STAT∷eRFP* reporter in developing embryos (from developmental stage 13 to 17). Maximum intensity projections of z-stack images that capture the surface of the embryo (25 μm) are shown (**Supplementary Video 2**). The surface signal was projected separately from the mid-section because it is much weaker than the signal from internal organs. STAT activity is initially detected at the tracheal pits (indicated by white arrowheads). As embryonic development continues, the dGFP signal disappears whereas the RFP signal remains. Meanwhile, STAT activity, revealed by *STAT∷edGFP* in the gonad (indicated by yellow arrow), also increases and decreases during the time of imaging. **c**, **d**, **e.** z-projection of the internal organs of a developing fly embryo expressing both *STAT∷edGFP* and *STAT∷eRFP* reporters (**Supplementary Video 1**). Fluorescent intensities of both GFP and RFP from three different positions, P1 (posterior spiracles and hindgut), P2 (pharynx), and P3 (proventriculus), are measured. Original and normalized total fluorescent signals from P1 (GFP1 and RFP1), P2 (GFP2 and RFP2), and P3 (GFP3 and RFP3) are plotted in **d** and **e**, respectively. **f.** Estimation of dGFP and RFP half-life (*T*_*GFP1/2*_ =2.1 hr, *T*_*RFP1/2*_ =18.5 hr, which represent the effects from both *T*_*m1/2*_ and *T*_*p1/2*_) using P1 (posterior spiracles and hindgut) signals from 6 different embryos. The degradation of dGFP is moderately deviated from the simplified model, suggesting that the substrate concentration may be comparable with the Michaelis-Menten (MM) constant. Therefore, the estimated half-life parameter is an average over time. Scale bar: **b.c**, 100 μm; **g.** 25 μm.

**Figure 4-figure supplement 1.**
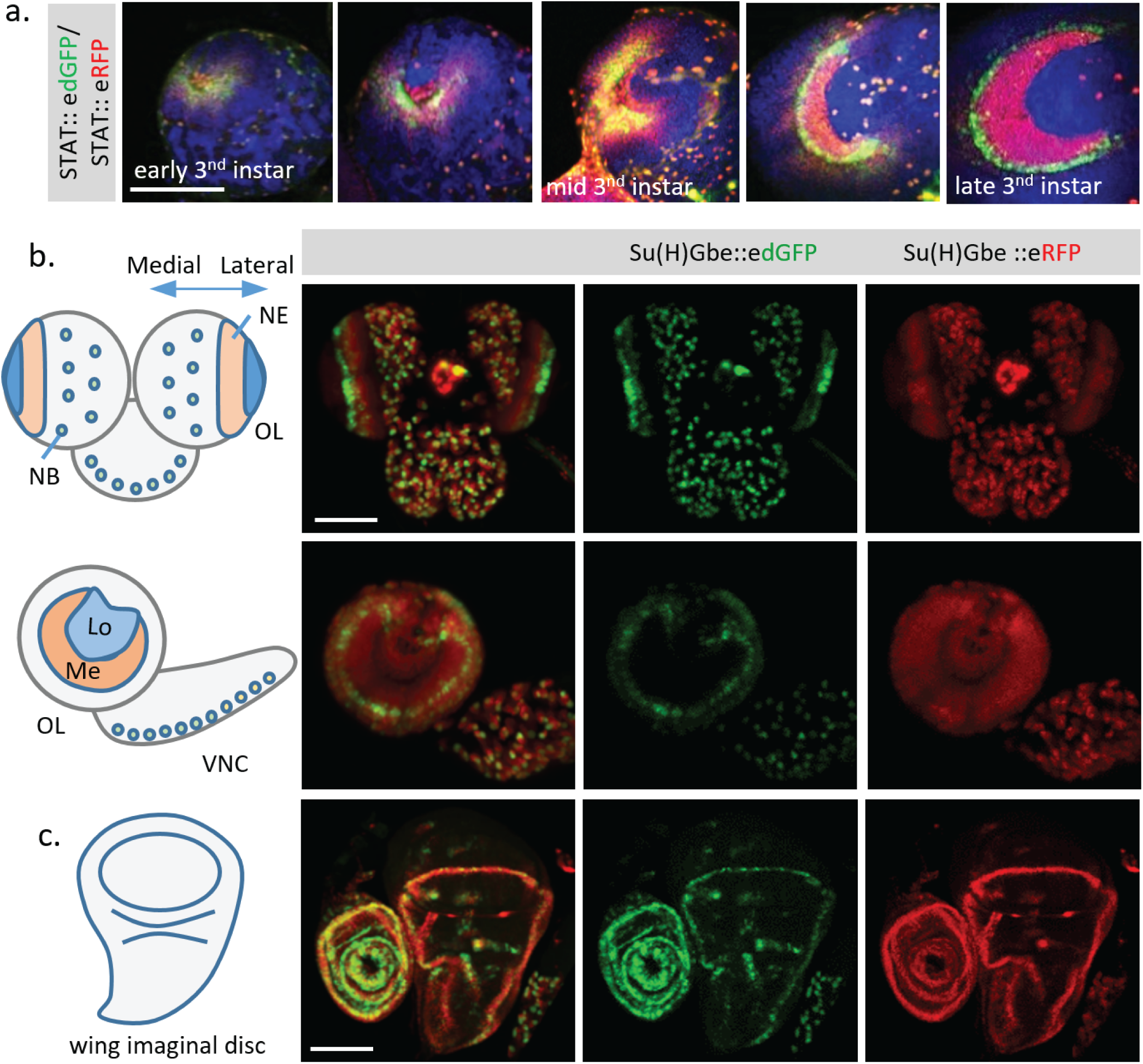
Additional dynamics of Notch and STAT activity revealed in different fixed tissues and stages. **a.** Change of STAT activity in 3^rd^ instar larval optic lobe (OL) at different developing stages. **b.** Activities of Notch signaling revealed by *Su(H)Gbe∷edGFP* and *Su(H)Gbe∷eRFP* in 3^rd^ instar larval brain. In the optic lobe (OL), the Notch signal is down-regulated by a “proneural wave” that sweeps across the neuroepithelium (NE) from the medial to the lateral regions, to trigger the NE to neuroblast (NB) transition. Consistent with this, a gradient change of Notch activity from red to green was observed in the NE region. Red signal (Notch low) located medially, and a band of green signal (Notch high) located at the lateral side of the OL, were observed, revealing the site of Notch pathway activation and progression of the proneural wave. Lo, lobula; Me, medulla; VNC, ventral nerve cord. **c.** Both dGFP and RFP reporters show activity in the wing margin region of the 3rd instar wing disc, with the GFP signal concentrated in a more defined cell population. Scale bar: **a**,**b**,**c.** 100 μm.

**Figure 4-figure supplement 2.**
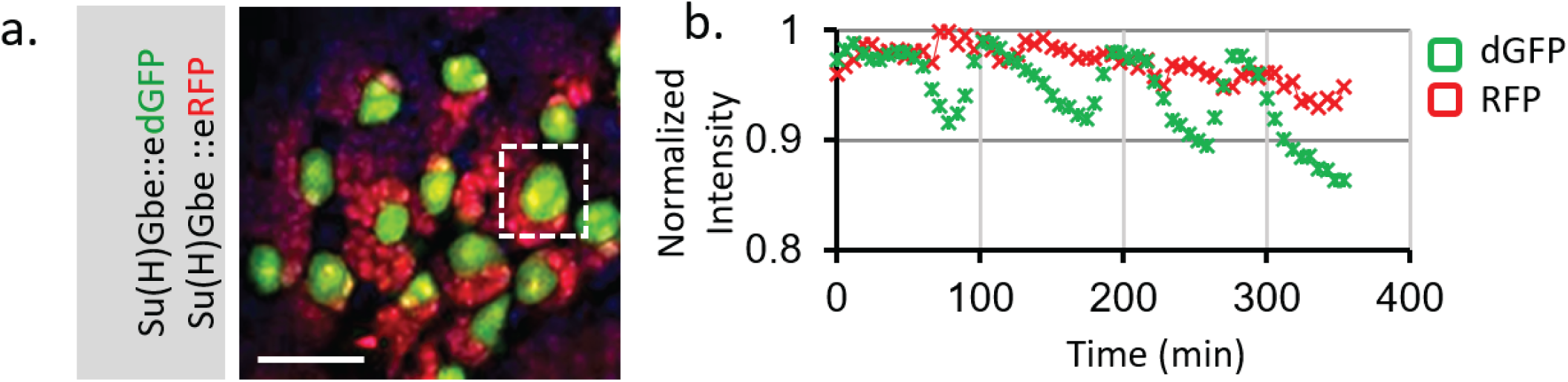
Live imaging of Notch activity in dividing Type I neuroblasts (NBs) in larval brain. **a.** Notch activity is monitored by both *Su(H)Gbe∷edGFP* and *Su(H)Gbe∷eRFP* reporters. Dissected 3^rd^ instar larval brain was cultured together with the larval fat body in Schneider’s medium and imaged every 5 min. A z-stack covering the entire NB cell body was taken. In the central brain region, the large neuroblasts (NBs) (green) undergo asymmetric cell division and create new NBs and much smaller progeny cells (red). The total signal from a single NB was measured using Bitplane Imaris software. The dGFP channel showed an oscillation with amplitude ∼8% of the average signal, whereas the RFP channel is generally flat. The change of dGFP is consistent with an asymmetric NB cell division event (∼ 1.5 hr) that produces a small progeny cell marked by RFP, suggesting that Notch activity may be affected during cell division. Scale bar: 25 μm.

**Figure 4-figure supplement 3.**
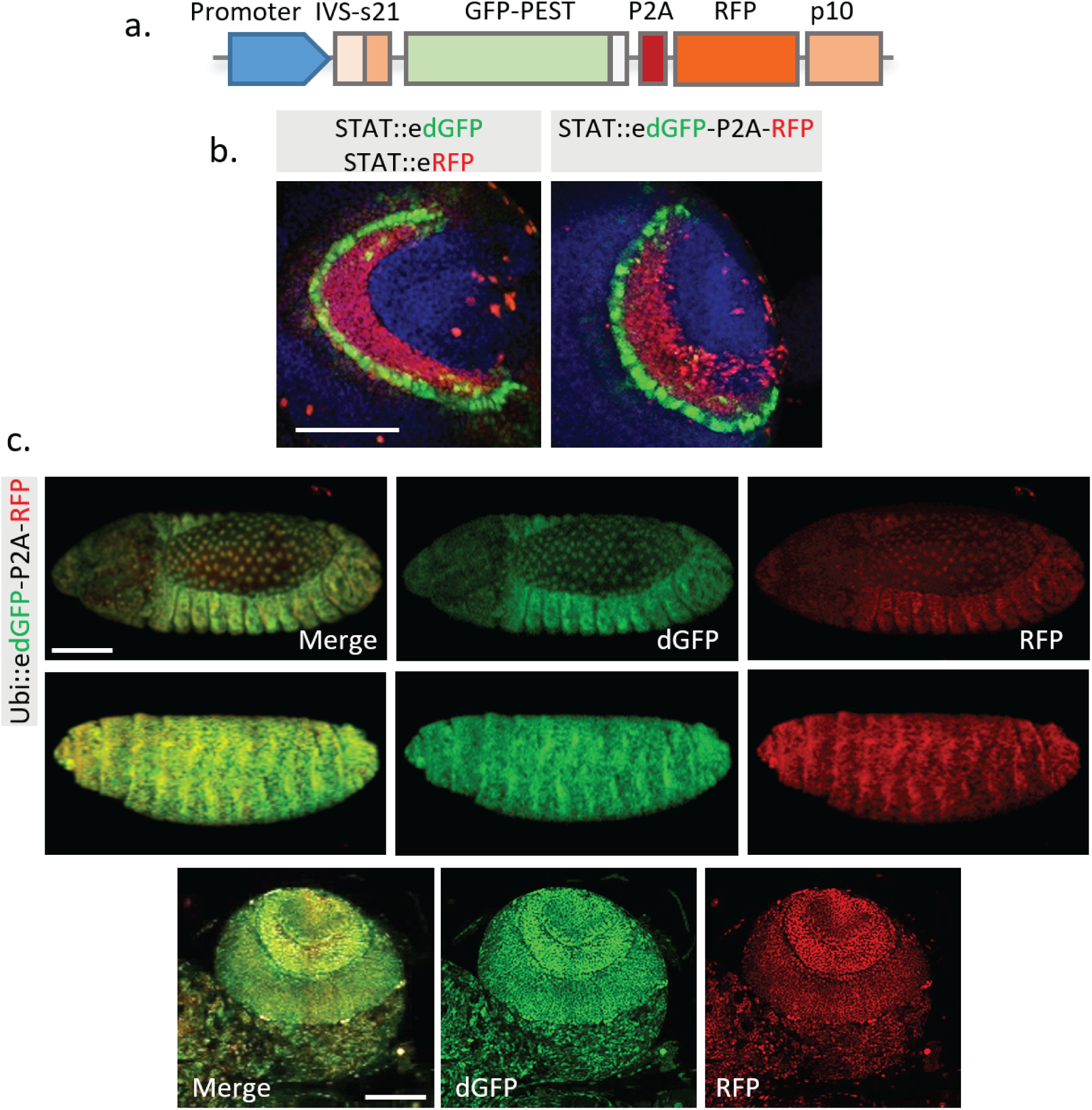
Combination of edGFP and RFP into a single construct using 2A peptide. **a.** Structure of the multicistronic reporter containing edGFP and RFP. **b.** Comparison between *STAT∷edGFP, STAT∷eRFP* and *STAT∷edGFP-P2A-RFP* in 3^rd^ instar larval brain. **c.** No significant change of ratio between dGFP compared to RFP was observed in the embryo and larval brain when both proteins are controlled by a constitutively active Ubiquitin promoter. Representative images from the stage 14 (top panel) and 16 (middle panel) fly embryos, and 3^rd^ instar larval optic lobe (bottom panel) are shown. Scale bar: **a**, 50 μm; **b**, 100 μm.

**Figure 5-figure supplement 1.**
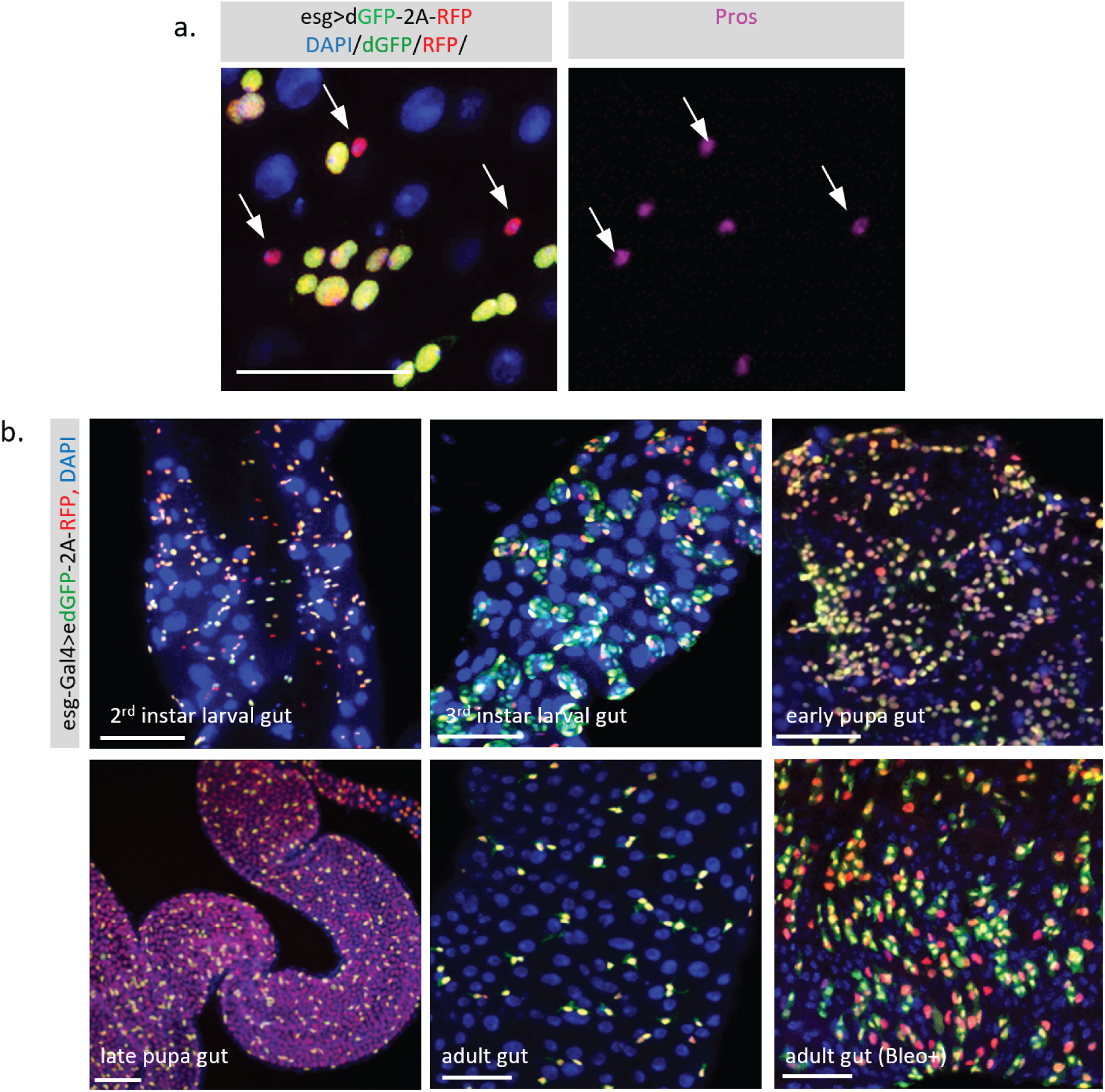
Dynamic activity of TransTimer in the fly intestine. **a.** The small cells with high RFP and low dGFP signals are positive for the enteroendocrine (EE) cell marker *Pros*, suggesting that these cells are differentiated cells that turn off *esg*. **b.** Patterns of the dynamic reporter driven by *esg-Gal4* in the *Drosophila* intestine at different developmental stages. In the 2nd instar, the adult midgut progenitors (AMPs) exist as individual cells and disperse throughout the intestine. Using TransTimer, we observed considerable variation in the dGFP/RFP ratio, indicating an unexpected heterogeneity in this cell population. In the 3rd instar intestine, the AMPs proliferate and form clusters called midgut imaginal islands (MIIs). Some AMPs with high Notch activity start to differentiate into peripheral cells (PCs), which surround the AMPs to create a transient stem cell niche. Consistent with this model, cells at the periphery of MII showed more red color, suggesting that these cells started to differentiate and lost their stemness. During the early pupal stage, the MIIs break down, while PCs and a large portion of AMPs differentiate into pupal intestine cells, which will be degraded at the end of the pupal stage. Consistent with this, many *esg*+ AMPs turn red as these cells start to differentiate. During metamorphosis, the larval intestine is completely turned over, and the late pupal intestine, which will become the future adult intestine, is newly generated from AMPs. Therefore, except for the ISCs that still maintain the dGFP signal, differentiated gut cells all show the RFP signal, suggesting that they were recently derived from *esg*+ AMPs. During the adult stage, *esg*+ ISCs and enteroblast (EBs) are mitotically inactive and predominately show a yellow color. However, in the presence of the damaging reagent Bleomycin, which triggers strong cell proliferation and differentiation, a significant separation between dGFP and RFP signal is observed. Scale bar: **a**, **b.** 50 μm.

**Figure 7-figure supplement 1.**
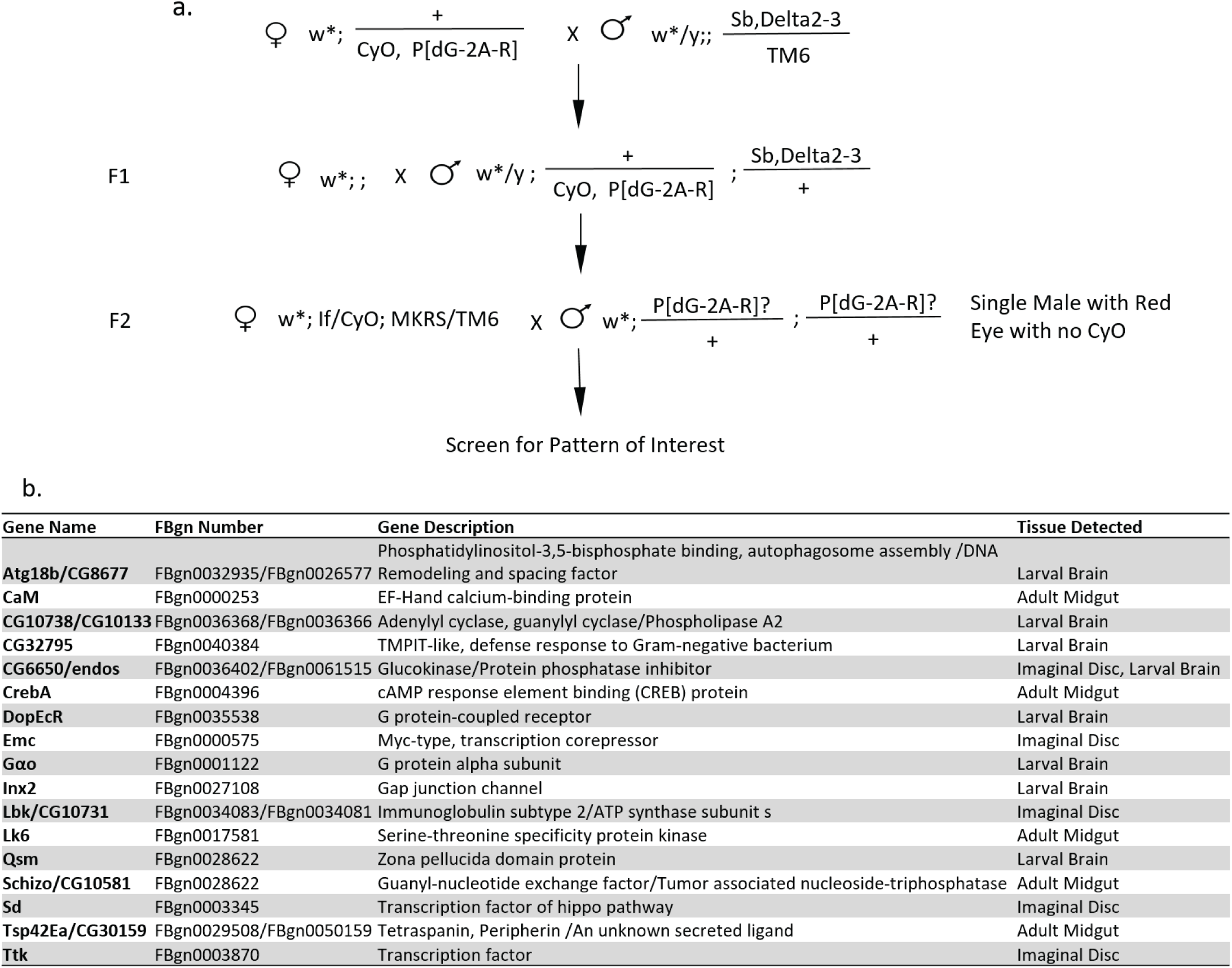
Enhancer trap screen for gene expression dynamics. **a.** Genetic crossing scheme for random insertions of a P-transposable element with TransTimer (*P[dG-2A-R]*) on the 2^nd^ & 3^rd^ chromosomes. Fly lines with interesting expression patterns were balanced and mapped using splinkerette PCR**. b.** List of identified enhancer trap lines with interesting dynamic expression patterns. The names of the genes with promoter regions located within 3 kb range of the reporter insertion sites are listed in the table. In most cases, the insertions showed a clear association with the promoter region of a specific gene. Representative images of the expression patterns are shown in **Supplementary data table 3**.

**Supplementary Video 1. Live imaging of developing fly embryo (mid projection).** Embryo was imaged at room temperature using a Zeiss Lightsheet Z1 microscope with a 20X (N.A. 1.0) lens. Z-stack images (2 μm between each slice) were acquired every 10 min. A maximal intensity z-projection of mid-section (120 μm) is shown in the video. Video was taken from stage 13 to stage 17 when the somatic muscles contract. Genotype of the sample is: w; STAT∷edGFP / STAT∷eRFP.

**Supplementary Video 2. Live imaging of developing fly embryo (surface projection).** Embryo was imaged at room temperature using a Zeiss Lightsheet Z1 microscope with a 20X (N.A. 1.0) lens. Z-stack images (2 μm between each slice) were acquired with 10 min time intervals. A maximal intensity z-projection of surface (25 μm) is shown. Video was taken from stage 13 to stage 17 before the somatic muscles contraction. Genotype of the sample is: w; STAT∷edGFP / STAT∷eRFP.

## References

1 Purvis, J. E. & Lahav, G. Encoding and decoding cellular information through signaling dynamics. Cell 152, 945–956, doi:10.1016/j.cell.2013.02.005 (2013).

2 Yosef, N. & Regev, A. Impulse control: temporal dynamics in gene transcription. Cell 144, 886–896, doi:10.1016/j.cell.2011.02.015 (2011).

3 Isomura, A. & Kageyama, R. Ultradian oscillations and pulses: coordinating cellular responses and cell fate decisions. Development 141, 3627–3636, doi:10.1242/dev.104497 (2014).

4 Doupe, D. P. & Perrimon, N. Visualizing and manipulating temporal signaling dynamics with fluorescence-based tools. Science signaling 7, re1, doi:10.1126/scisignal.2005077 (2014).

5 Flores, G. V. et al. Combinatorial signaling in the specification of unique cell fates. Cell 103, 75–85 (2000).

6 Tsuda, L., Nagaraj, R., Zipursky, S. L. & Banerjee, U. An EGFR/Ebi/Sno pathway promotes delta expression by inactivating Su(H)/SMRTER repression during inductive notch signaling. Cell 110, 625–637 (2002).

7 Bending, D. et al. A timer for analyzing temporally dynamic changes in transcription during differentiation in vivo. The Journal of cell biology 217, 2931–2950, doi:10.1083/jcb.201711048 (2018).

8 Subach, F. V. et al. Monomeric fluorescent timers that change color from blue to red report on cellular trafficking. Nature chemical biology 5, 118–126, doi:10.1038/nchembio.138 (2009).

9 Li, X. et al. Generation of destabilized green fluorescent protein as a transcription reporter. The Journal of biological chemistry 273, 34970–34975 (1998).

10 Rogers, S., Wells, R. & Rechsteiner, M. Amino acid sequences common to rapidly degraded proteins: the PEST hypothesis. Science 234, 364–368 (1986).

11 Pedelacq, J. D., Cabantous, S., Tran, T., Terwilliger, T. C. & Waldo, G. S. Engineering and characterization of a superfolder green fluorescent protein. Nature biotechnology 24, 79–88, doi:10.1038/nbt1172 (2006).

12 Gordon, A. et al. Single-cell quantification of molecules and rates using open-source microscope-based cytometry. Nat Methods 4, 175–181, doi:10.1038/nmeth1008 (2007).

13 Baker, K. E. & Parker, R. Conventional 3’ end formation is not required for NMD substrate recognition in Saccharomyces cerevisiae. Rna 12, 1441–1445, doi:10.1261/rna.92706 (2006).

14 Sacchetti, A., El Sewedy, T., Nasr, A. F. & Alberti, S. Efficient GFP mutations profoundly affect mRNA transcription and translation rates. FEBS Lett 492, 151–155 (2001).

15 Houser, J. R. et al. An improved short-lived fluorescent protein transcriptional reporter for Saccharomyces cerevisiae. Yeast 29, 519–530, doi:10.1002/yea.2932 (2012).

16 Voon, D. C. et al. Use of mRNA-and protein-destabilizing elements to develop a highly responsive reporter system. Nucleic Acids Res 33, e27, doi:10.1093/nar/gni030 (2005).

17 Venken, K. J. et al. MiMIC: a highly versatile transposon insertion resource for engineering Drosophila melanogaster genes. Nature methods 8, 737–743 (2011).

18 Bach, E. A. et al. GFP reporters detect the activation of the Drosophila JAK/STAT pathway in vivo. Gene expression patterns : GEP 7, 323–331, doi:10.1016/j.modgep.2006.08.003 (2007).

19 Furriols, M. & Bray, S. A model Notch response element detects Suppressor of Hairless-dependent molecular switch. Current biology : CB 11, 60–64 (2001).

20 Cranfill, P. J. et al. Quantitative assessment of fluorescent proteins. Nature methods 13, 557–562, doi:10.1038/nmeth.3891 (2016).

21 Shearin, H. K., Macdonald, I. S., Spector, L. P. & Stowers, R. S. Hexameric GFP and mCherry reporters for the Drosophila GAL4, Q, and LexA transcription systems. Genetics 196, 951–960, doi:10.1534/genetics.113.161141 (2014).

22 Genove, G., Glick, B. S. & Barth, A. L. Brighter reporter genes from multimerized fluorescent proteins. BioTechniques 39, 814, 816, 818 passim (2005).

23 Pfeiffer, B. D., Truman, J. W. & Rubin, G. M. Using translational enhancers to increase transgene expression in Drosophila. Proceedings of the National Academy of Sciences of the United States of America 109, 6626–6631, doi:10.1073/pnas.1204520109 (2012).

24 Pfeiffer, B. D. et al. Refinement of tools for targeted gene expression in Drosophila. Genetics 186, 735–755, doi:10.1534/genetics.110.119917 (2010).

25 Suzuki, T. et al. Performance of expression vector, pTD1, in insect cell-free translation system. Journal of bioscience and bioengineering 102, 69–71, doi:10.1263/jbb.102.69 (2006).

26 van Oers, M. M., Vlak, J. M., Voorma, H. O. & Thomas, A. A. Role of the 3’ untranslated region of baculovirus p10 mRNA in high-level expression of foreign genes. The Journal of general virology 80 (Pt 8), 2253–2262, doi:10.1099/0022-1317-80-8-2253 (1999).

27 Groth, A. C., Fish, M., Nusse, R. & Calos, M. P. Construction of transgenic Drosophila by using the site-specific integrase from phage phiC31. Genetics 166, 1775–1782 (2004).

28 Rodrigues, A. B. et al. Activated STAT regulates growth and induces competitive interactions independently of Myc, Yorkie, Wingless and ribosome biogenesis. Development 139, 4051–4061, doi:10.1242/dev.076760 (2012).

29 Johansen, K. A., Iwaki, D. D. & Lengyel, J. A. Localized JAK/STAT signaling is required for oriented cell rearrangement in a tubular epithelium. Development 130, 135–145 (2003).

30 Johnson, A. N., Mokalled, M. H., Haden, T. N. & Olson, E. N. JAK/Stat signaling regulates heart precursor diversification in Drosophila. Development 138, 4627–4638, doi:10.1242/dev.071464 (2011).

31 Khmelinskii, A. et al. Tandem fluorescent protein timers for in vivo analysis of protein dynamics. Nature biotechnology 30, 708–714, doi:10.1038/nbt.2281 (2012).

32 Merzlyak, E. M. et al. Bright monomeric red fluorescent protein with an extended fluorescence lifetime. Nat Methods 4, 555–557, doi:10.1038/nmeth1062 (2007).

33 Gjetting, K. S., Ytting, C. K., Schulz, A. & Fuglsang, A. T. Live imaging of intra-and extracellular pH in plants using pHusion, a novel genetically encoded biosensor. Journal of experimental botany 63, 3207–3218, doi:10.1093/jxb/ers040 (2012).

34 Zhou, C. et al. Monitoring autophagic flux by an improved tandem fluorescent-tagged LC3 (mTagRFP-mWasabi-LC3) reveals that high-dose rapamycin impairs autophagic flux in cancer cells. Autophagy 8, 1215–1226, doi:10.4161/auto.20284 (2012).

35 Koldenkova, V. P., Matsuda, T. & Nagai, T. MagIC, a genetically encoded fluorescent indicator for monitoring cellular Mg2+ using a non-Forster resonance energy transfer ratiometric imaging approach. Journal of biomedical optics 20, 101203, doi:10.1117/1.JBO.20.10.101203 (2015).

36 Yasugi, T., Umetsu, D., Murakami, S., Sato, M. & Tabata, T. Drosophila optic lobe neuroblasts triggered by a wave of proneural gene expression that is negatively regulated by JAK/STAT. Development 135, 1471–1480, doi:10.1242/dev.019117 (2008).

37 Wang, H. et al. Aurora-A acts as a tumor suppressor and regulates self-renewal of Drosophila neuroblasts. Genes & development 20, 3453–3463, doi:10.1101/gad.1487506 (2006).

38 Szymczak, A. L. & Vignali, D. A. Development of 2A peptide-based strategies in the design of multicistronic vectors. Expert opinion on biological therapy 5, 627–638, doi:10.1517/14712598.5.5.627 (2005).

39 Brand, A. H. & Perrimon, N. Targeted gene expression as a means of altering cell fates and generating dominant phenotypes. Development 118, 401–415 (1993).

40 Jenett, A. et al. A GAL4-driver line resource for Drosophila neurobiology. Cell reports 2, 991– 1001, doi:10.1016/j.celrep.2012.09.011 (2012).

41 Lee, P. T. et al. A gene-specific T2A-GAL4 library for Drosophila. eLife 7, doi:10.7554/eLife.35574 (2018).

42 Micchelli, C. A. & Perrimon, N. Evidence that stem cells reside in the adult Drosophila midgut epithelium. Nature 439, 475–479, doi:10.1038/nature04371 (2006).

43 Ohlstein, B. & Spradling, A. Multipotent Drosophila intestinal stem cells specify daughter cell fates by differential notch signaling. Science 315, 988–992, doi:10.1126/science.1136606 (2007).

44 Amcheslavsky, A., Jiang, J. & Ip, Y. T. Tissue damage-induced intestinal stem cell division in Drosophila. Cell stem cell 4, 49–61, doi:10.1016/j.stem.2008.10.016 (2009).

45 Meacham, C. E. & Morrison, S. J. Tumour heterogeneity and cancer cell plasticity. Nature 501, 328–337, doi:10.1038/nature12624 (2013).

46 Jiang, H. et al. Cytokine/Jak/Stat signaling mediates regeneration and homeostasis in the Drosophila midgut. Cell 137, 1343–1355, doi:10.1016/j.cell.2009.05.014 (2009).

47 Marianes, A. & Spradling, A. C. Physiological and stem cell compartmentalization within the Drosophila midgut. eLife 2, e00886, doi:10.7554/eLife.00886 (2013).

48 He, L., Si, G., Huang, J., Samuel, A. D. T. & Perrimon, N. Mechanical regulation of stem-cell differentiation by the stretch-activated Piezo channel. Nature 555, 103–106, doi:10.1038/nature25744 (2018).

49 Hao, D. et al. AC3-33, a novel secretory protein, inhibits Elk1 transcriptional activity via ERK pathway. Molecular biology reports 38, 1375–1382, doi:10.1007/s11033-010-0240-x (2011).

50 Yagi, R., Mayer, F. & Basler, K. Refined LexA transactivators and their use in combination with the Drosophila Gal4 system. Proceedings of the National Academy of Sciences of the United States of America 107, 16166–16171, doi:10.1073/pnas.1005957107 (2010).

51 Nern, A., Pfeiffer, B. D., Svoboda, K. & Rubin, G. M. Multiple new site-specific recombinases for use in manipulating animal genomes. Proceedings of the National Academy of Sciences of the United States of America 108, 14198–14203, doi:10.1073/pnas.1111704108 (2011).

52 Zhu, S., Barshow, S., Wildonger, J., Jan, L. Y. & Jan, Y. N. Ets transcription factor Pointed promotes the generation of intermediate neural progenitors in Drosophila larval brains. Proceedings of the National Academy of Sciences of the United States of America 108, 20615– 20620, doi:10.1073/pnas.1118595109 (2011).

53 Amcheslavsky, A. et al. Enteroendocrine cells support intestinal stem-cell-mediated homeostasis in Drosophila. Cell reports 9, 32–39, doi:10.1016/j.celrep.2014.08.052 (2014).

54 Eaton, S. L. et al. A guide to modern quantitative fluorescent western blotting with troubleshooting strategies. Journal of visualized experiments : JoVE, e52099, doi:10.3791/52099 (2014).

55 Rubin, G. M. & Spradling, A. C. Genetic transformation of Drosophila with transposable element vectors. Science 218, 348–353 (1982).

56 Horn, C. et al. Splinkerette PCR for more efficient characterization of gene trap events. Nature genetics 39, 933–934, doi:10.1038/ng0807-933 (2007).

57 Potter, C. J. & Luo, L. Splinkerette PCR for mapping transposable elements in Drosophila. PloS one 5, e10168, doi:10.1371/journal.pone.0010168 (2010).

58 Wang, X., Errede, B. & Elston, T. C. Mathematical analysis and quantification of fluorescent proteins as transcriptional reporters. Biophysical journal 94, 2017–2026, doi:10.1529/biophysj.107.122200 (2008).

59 Tomer, R., Khairy, K., Amat, F. & Keller, P. J. Quantitative high-speed imaging of entire developing embryos with simultaneous multiview light-sheet microscopy. Nature methods 9, 755–763, doi:10.1038/nmeth.2062 (2012).

60 Lerit, D. A., Plevock, K. M. & Rusan, N. M. Live imaging of Drosophila larval neuroblasts. Journal of visualized experiments : JoVE, doi:10.3791/51756 (2014).

61 Lemon, W. C. et al. Whole-central nervous system functional imaging in larval Drosophila. Nature communications 6, 7924, doi:10.1038/ncomms8924 (2015).

